# Hypomorphic *Lig4* gene mutation in mice predisposes to Th1-skewing intestinal inflammation

**DOI:** 10.1101/2025.05.14.654009

**Authors:** Yusuke Yamashita, Hideki Kosako, Takashi Kato, Izumi Sasaki, Sadahiro Iwabuchi, Tadashi Okamura, Misato Tane, Shotaro Tabata, Kazutaka Nakashima, Ken Tanaka, Kazunori Shiraishi, Yuki Uchihara, Daisuke Okuzaki, Atsushi Shibata, Tsunehiro Mizushima, Hiroaki Hemmi, Nobuo Kanazawa, Seiji Kodama, Kouichi Ohshima, Shinichi Hashimoto, Yoshio Fujitani, Takashi Sonoki, Shinobu Tamura, Tsuneyasu Kaisho

## Abstract

Dysregulation of DNA double-strand break (DSB) repair leads to adaptive immunodeficiency, whereas the remaining lymphocytes are aberrantly activated and provoke inflammations. However, no model mice were available to consistently manifest inflammation under defective DSB repair. We generated mutant mice carrying a missense mutation p.W447C in the gene encoding DNA ligase IV (LIG4), critical for DSB repair. *Lig4^W447C/W447C^* mice showed growth retardation and severe intestinal inflammations under adaptive immunodeficiency. The inflammations were featured by marked infiltration of T helper type 1 (Th1) cells and macrophages and was dependent on lymphocytes. When *Ifng* was deleted, Th2 and Th17 instead of Th1 cells drove the inflammations. *Lig4^W447C/W447C^* mice showed expansion of oligoclonal T cells with T cell receptor α repertoire skewed towards more proximal 3’ V and 5’ J gene segments. Thus, our novel hypomorphic *Lig4* mutant mice show that defective DSB repair leads to Th1-dependent intestinal inflammations under severe adaptive immunodeficiency.

## Introduction

In mammals, DNAs are damaged to generate DNA double-strand breaks (DSBs) not only upon exposure to ionizing radiation or ultraviolet, but also in the physiological development of nervous and immune systems. Non-homologous end joining (NHEJ) is a critical pathway to repair the DSBs ^1, 2, 3^. In lymphocytes, NHEJ is indispensable for productive V(D)J recombination of T cell receptor (TCR) and immunoglobulin (Ig) genes. NHEJ is mediated by a series of reactions, in which there is involvement of DNA-dependent protein kinase catalytic subunit (DNA-PKcs), the KU70/80 heterodimer, Artemis, Cernunnos (also known as Xrcc4-like factor, XLF), DNA ligase IV (LIGIV), and X-ray repair cross-complementing protein 4 (XRCC4) ^1, 2, 3^. In humans, defects in any of these components can lead to profound acquired immunodeficiency by impairment of the V(D)J recombination, increased radiosensitivity and neurodevelopmental delay ^4^.

LIGIV binds to DSB ends and acts as the ligase with XRCC4 in the final end-joining step of NHEJ ^5, 6^. Hypomorphic variants in the human *LIG4* gene underlie LIG4 syndrome, a rare autosomal recessive disorder characterized by growth disturbance, neurodevelopmental delay, increased radiosensitivity, and adaptive immunodeficiency ^5, 6, 7^. Patients with LIG4 syndrome have also shown a predisposition to malignancies including leukemia and lymphoma ^8, 9, 10, 11^. Paradoxically, because of dysregulated activation of adaptive immunity, they occasionally exhibit inflammatory disorders such as inflammatory bowel disease and Bechet diseases ^12, 13, 14^. Although LIG4 syndrome exhibits phenotypic heterogeneity, little is known about how the variants of the human *LIG4* gene contribute to diverse manifestations such as inflammatory conditions.

In mice, *Lig4* deficiency causes embryonic lethality due to impaired neuronal development ^15, 16^. Two kinds of mutant mice carrying a hypomorphic *Lig4* variant in the enzymatic domain have been generated to date, and homozygous mutant mice showed neurodevelopmental delay from birth as well as adaptive immunodeficiency ^17, 18, 19, 20, 21^. In mice homozygous for a *Lig4* R278H mutation (*Lig4^R278H/R278H^* mice) derived from a patient with LIG4 syndrome, thymic T-cell lymphoma was observed at high frequencies ^19^. Moreover, *Lig4^Y288C/Y288C^* mice, which were generated by *N*-ethyl-*N*-nitrosourea (ENU), also exhibited a high incidence of lymphoid neoplasms that originate from the thymus ^20^. LIG4 syndrome manifestations were thus well recapitulated in the mutant mice. However, none of these mutant mice show any apparent signs of inflammation.

We have previously described a patient with LIG4 syndrome that was carrying a compound heterozygous variant in the *LIG4* gene. One variant was a nonsense variant, p.E413X, the other one was a missense variant, p.W447C, found in the well-conserved amino acid among species ^22^. In the current study, we generated mutant mice carrying this missense variant. *Lig4^W447C/W447C^*mice manifested not only neuronal and developmental defects, but also lymphocyte-dependent colitis with acquired immunodeficiency. The colitis was featured by infiltration of type 1 helper T (Th1) cells and macrophages and lack of B cells. *Lig4^W447C/W447C^*mice are unique immunodeficient mice manifesting Th1 cell-driven inflammatory disorders.

## Results

### *Lig4^W447C/W447C^* mice showed microcephaly, growth retardation, and radiosensitivity

To clarify the pathological roles of a missense variant (p.W447C), we introduced the variant in mice by the CRISPR-Cas9 system. *Lig4^W447C/+^* mice were born and appeared healthy. After crossing between *Lig4^W447C/+^* mice, *Lig4^W447C/W447C^*mice were born with expected Mendelian inheritance but showed short stature and low birth weight **(Extended Data Fig. 1a)**. The mice also exhibited growth retardation and microcephaly **(Fig. 1a)**. These manifestations are hallmarks of the LIG4 syndrome ^5, 6, 7^. We have then investigated DSBs by γH2AX foci formation in mouse embryonic fibroblasts (MEFs). The formation was comparable between *Lig4^+/+^* and *Lig4^W447C/+^*MEFs. In *Lig4^W447C/W447C^* MEFs, more γH2AX foci was detected not only before, but also upon irradiation than in *Lig4^W447C/+^* MEFs **(Fig. 1b and Extended Data Fig. 1b)**. Thus, *Lig4^W447C/W447C^* mice showed growth retardation, microcephaly, and defects in DNA damage repair.

**Figure 1.**
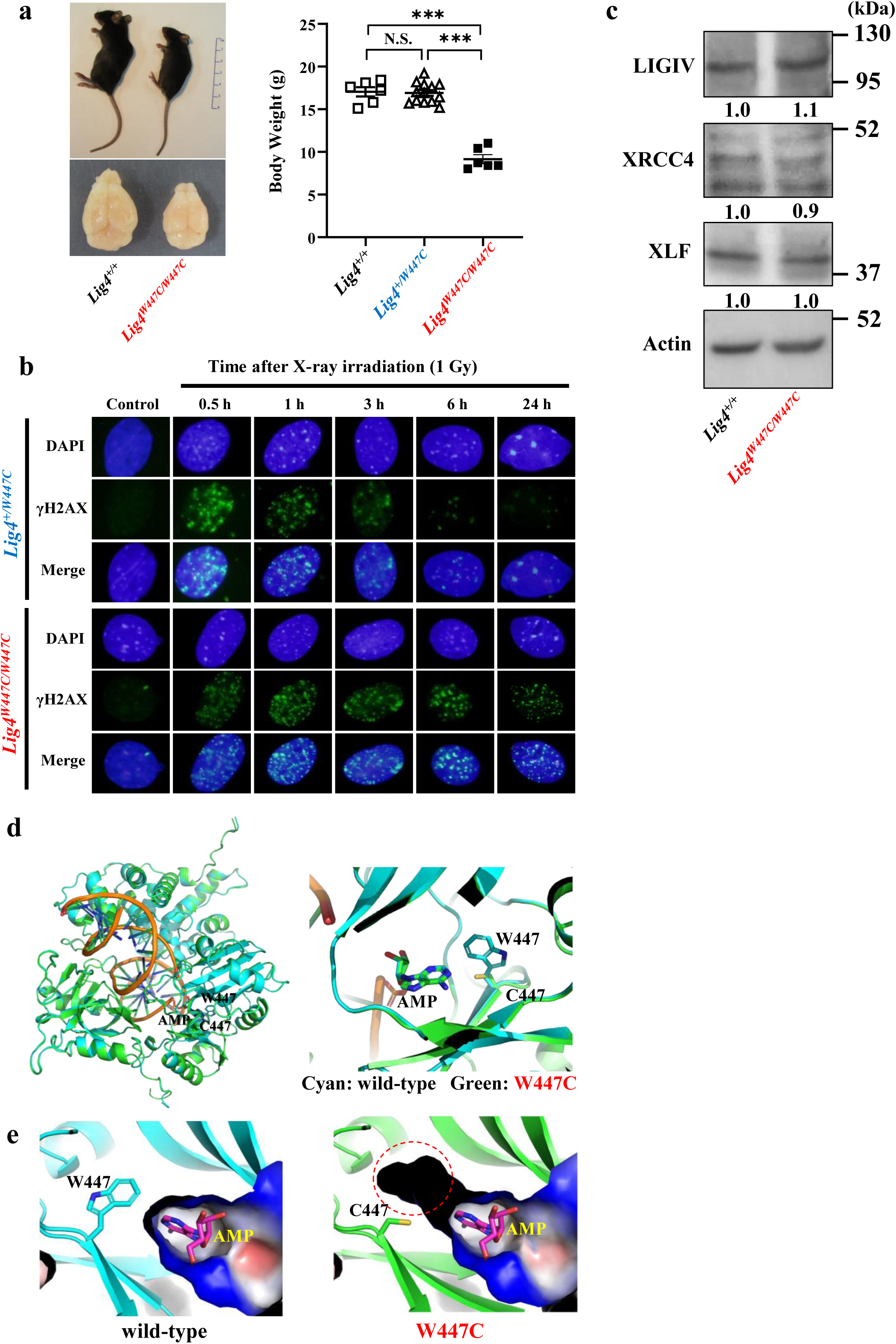
*Lig4^W447C/W447C^* mice displayed typical LIG4 syndrome features. (a) The appearance and brain of six-week-old *Lig4*^+/+^ and *Lig4^W447C/W447C^* littermates. The graph (right) depicts the weight of six-week-old female mice (*Lig4*^+/+^ [*n* = 6], *Lig4^+/W447C^* [*n* = 13] and *Lig4^W447C/W447C^*[*n* = 6]). *P*-values were calculated using one-way analysis of variance (ANOVA) followed by Tukey’s post hoc multiple comparison test. (b) Immunofluorescence of γH2AX foci in the nuclei of *Lig4*^+/W447C^ and *Lig4^W447C/W447C^* mouse embryonic fibroblasts (MEFs) at each representative time after 1 Gy irradiation. (c) Western blots of LIGIV, XRCC4, XLF and β-actin protein expressions on *Lig4^+/+^*and *Lig4^W447C/W447C^* MEFs. Numbers below the representative bands indicate the ratio of signal intensity relative to *Lig4^+/+^* after normalization to β-actin, as measured using Image J software. (d) Mouse LIGIV protein structures in wild-type and W447C-mutant mice (left panel). The overall LIG IV protein structure, with DNA-binding and enzymatic domains (residues 1–620), is depicted as a ribbon model (right panel). W447 and C447 binding to adenosine monophosphate (AMP) are shown in stick representation (cyan and green, respectively). (e) Structural comparison between W447 (left panel, cyan) and C447 (right panel, green). AlphaFold3-predicted side chains related to AMP-binding are substantially different between them. Consequently, W447C lead to the enlargement of AMP-binding pocket (dotted red circle in right panel), suggesting the destabilization of AMP-binding site during DNA repair processes (black region). The data are presented as the mean ± SEM.

Expression of LIGIV protein in *Lig4^W447C/W447C^* MEFs was at comparable levels to that in wild-type MEFs, indicating that the mutation does not affect the amounts of LIGIV protein **(Fig. 1c)**. In addition, the expression levels of XRCC4 and XLF, which assist LIGIV in its DNA repair function, were also unchanged between *Lig4^W447C/W447C^*and wild-type MEFs **(Fig. 1c)**. We therefore performed *in silico* analysis on wild-type and W447C mutant of LIGIV. W447 was localized in face to adenosine monophosphate (AMP) in the enzymatic domain. The W447C mutant did not affect overall structures, but it resulted in a decrease of the AMP-binding interface area (wild-type 2.66 Å^2^, W447C mutant 1.98 Å^2^) and enlargement of the AMP-binding pocket mainly due to lack of an indole ring **(Fig. 1d, e)** ^23^. We also analyzed the R278H and Y288C mutants of LIGIV **(Extended Data Fig. 2a, b)**, but neither mutant altered overall structures of LIGIV. Y288 did not face to AMP, while R278 was in contact with AMP and the R278H mutant decreased the interface area (wild-type 27.84 Å^2^, W447C mutant 18.46 Å^2^), but did not significantly affect the structure of the AMP-binding pocket.

### *Lig4^W447C/W447C^* mice exhibited severe adaptive immunodeficiency

We then analyzed the phenotype of lymphocytes in *Lig4^+/+^* and *Lig4^W447C/W447C^* mice. *Lig4^W447C/W447C^* mice had half the number of bone marrow (BM) cells of *Lig4^+/+^* mice **(Fig. 2a, b)**. In the BM of *Lig4^W447C/W447C^* mice, B220^+^CD43^+^IgM^−^ pro-B cells were decreased and there was marked reduction in B220^+^CD43^−^IgM^−^ pre-B cells, B220^low^CD43^−^IgM^+^ immature B cells and B220^high^CD43^−^IgM^+^ mature B cells **(Fig. 2c, d)**. Meanwhile, the proportion of pro-B cells among B220^+^ cells was significantly increased, indicating incomplete block at the pro-B cell stage, which was consistent with impaired rearrangement of the IgH locus **(Fig. 2d)**. In the spleens of *Lig4^W447C/W447C^* mice, follicular formation was severely impaired and the numbers of splenocytes and B220+ cells were severely decreased. **(Fig. 2e-h)**. Furthermore, in *Lig4^W447C/W447C^*mice, serum IgM, IgG1, and IgA levels were severely decreased **(Fig. 2i)**. Notably, serum IgE levels remained low, in similar to *Lig4^+/+^* mice **(Fig. 2i)**. In addition, the serum levels of anti-dsDNA antibody in *Lig4^W447C/W447C^* mice were lower than those of *Lig4*^+/+^ mice **(Extended Data Fig. 3).** Thus, B cell generation was defective in *Lig^4W447C/W447C^* mice.

**Figure 2.**
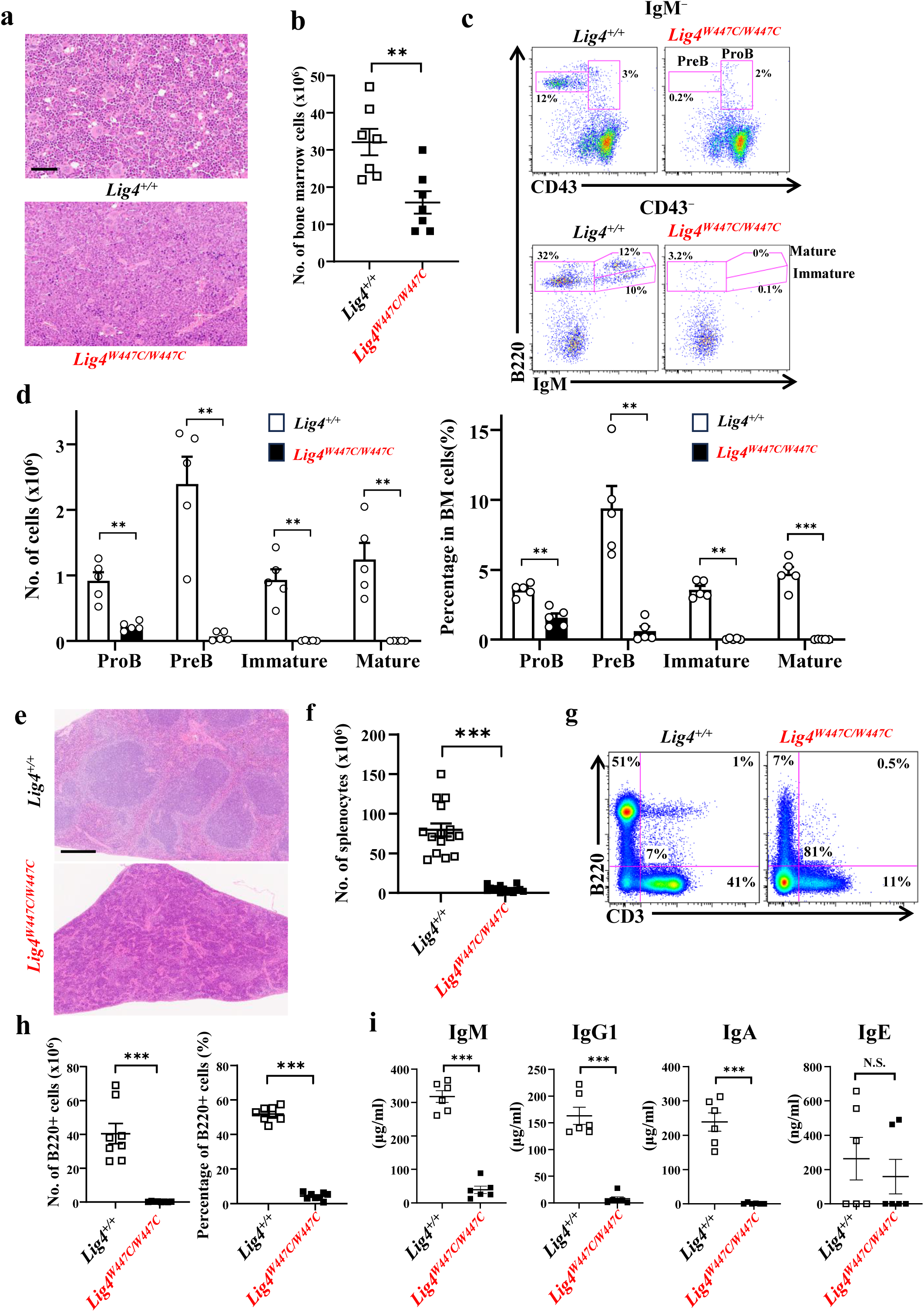
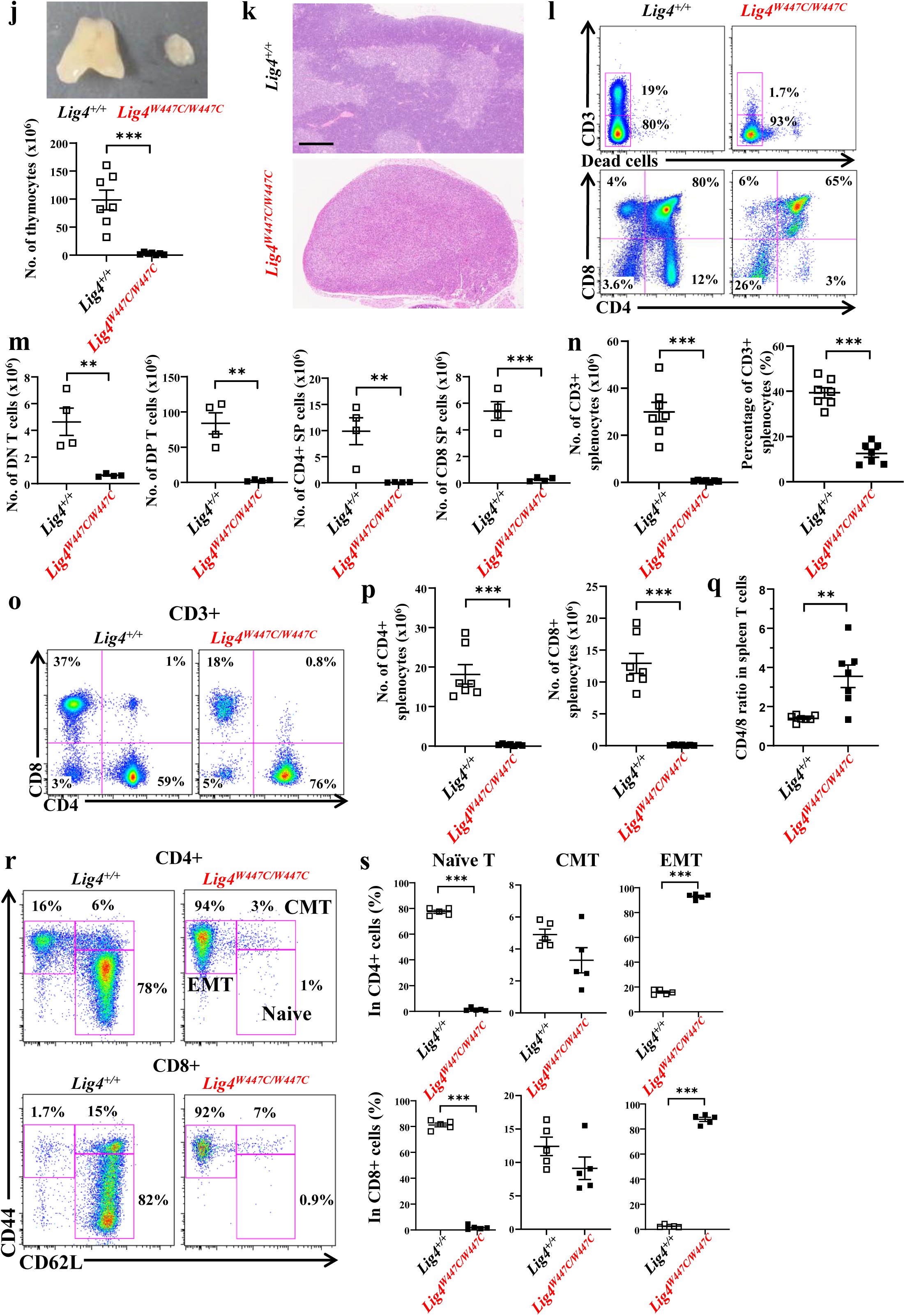
B– and T-cell differentiation was impaired in *Lig4^W447C/W447C^* mice. (a) Hematoxylin and eosin (H&E) staining revealed hypocellular marrow in bone marrow of *Lig4^W447C/W447C^* mouse compared with those of *Lig4^W447C/W447C^* mouse. Scale bar represents 60 μm. (b) The number of bone marrow (BM) cells in *Lig4^W447C/W447C^* mice was significantly lower than in *Lig4^+/+^* mice (*Lig4^+/+^* [*n* = 7] and in *Lig4^W447C/W447C^* mice [*n* = 7]; 7–23 weeks). (c) Dot plot analysis of BM cells labeled with B220, IgM, and CD43 antibodies. (d) The number and percentage of B cells in BM in *Lig4*^+/+^ and *Lig4^W447C/W447C^* mice (mean ± standard error of the mean [SEM], *Lig4^+/+^* [*n* = 5] and *Lig4^W447C/W447C^*[*n* = 5]; 8–30 weeks) (e) H&E staining of spleen from *Lig4^+/+^*and *Lig4^W447C/W447C^* mice. Scale bar represents 300 μm. (f) The graph depicts the number of splenocytes in *Lig4^+/+^* and *Lig4^W447C/W447C^* mice (*Lig4^+/+^*[*n* = 15] and *Lig4^W447C/W447C^*[*n* = 15]; 7–20 weeks). (g) Fluorescence-activated cell sorting (FACS) analysis of splenocytes labeled with CD3 and B220 antibodies from *Lig4^+/+^*and *Lig4^W447C/W447C^* mice. (h) The graphs depict the absolute number and percentage of B220+ splenocytes analyzed by FACS in *Lig4^+/+^* and *Lig4^W447C/W447C^* mice (*Lig4^+/+^* [n = 8] and *Lig4^W447C/W447C^* [n = 8]; 7–20 weeks). (i) Serum immunoglobulin (IgM, IgG1, IgA, and IgE) levels in *Lig4^+/+^* and *Lig4^W447C/W447C^* mice (*Lig4^+/+^* [*n* = 6] and *Lig4^W447C/W447C^*[*n* = 6]; 9–12 weeks). (j) Thymic hypoplasia in a six-week-old *Lig4^W447C/W447C^* mouse. The graph (below) depicts the number of thymocytes in *Lig4*^+/+^ and *Lig4^W447C/W447C^* mice (*Lig4^+/+^* [*n* = 7] and *Lig4^W447C/W447C^* [*n* = 7]; 8–23 weeks). (k) H&E staining of thymus from *Lig4^+/+^* and *Lig4^W447C/W447C^*mice. Scale bar represents 300 μm. (l) FACS analysis of thymocytes labeled with CD3, CD4, and CD8 antibodies from *Lig4*^+/+^ and *Lig4^W447C/W447C^*mice. (m) The numbers dynamics of thymic T cells at various developmental stages, including double negative, double positive, and single positive (SP) in *Lig4^+/+^* and *Lig4^W447C/W447C^* mice (*Lig4^+/+^* [n = 4] and *Lig4^W447C/W447C^* [n = 4]; 7–16 weeks). (n) The graph depicts the number and percentage of CD3+ splenocytes in *Lig4^+/+^* and *Lig4^W447C/W447C^* mice (*Lig4^+/+^* [*n* = 7] and *Lig4^W447C/W447C^* [*n* = 7]; 8–11 weeks). (o) FACS analysis of splenocytes labeled with CD3, CD4 and CD8 antibodies from *Lig4^+/+^* and *Lig4^W447C/W447C^* mice. (p) The absolute number of CD4+ and CD8+ splenocytes in *Lig4^+/+^* and *Lig4^W447C/W447C^*mice (*Lig4^+/+^* [*n* = 7] and *Lig4^W447C/W447C^*[*n* = 7]; 8–32 weeks). (q) The graph depicts the CD4/CD8 ratio of CD3+ spleen T cells in *Lig4^+/+^* mice versus *Lig4^W447C/W447C^* mice (*Lig4^+/+^* [*n* = 7] and *Lig4^W447C/W447C^* [*n* = 7]; 8–32 weeks). (r) FACS analysis of CD44 and CD62L expression in CD4^+^ or CD8^+^ T cells from the spleens of *Lig4^+/+^* and *Lig4^W447C/W447C^* mice. (s) T-cell subsets, naive, central memory (CMT), and effector memory (EMT) percentages in the spleens of *Lig4^+/+^* and *Lig4^W447C/W447C^* mice (*Lig4^+/+^* [*n* = 5] and *Lig4^W447C/W447C^* [*n* = 5]; 9–10 weeks). The data are presented as the mean ± SEM.

In *Lig4^W447C/W447C^* mice, the thymus was very small **(Fig. 2j)** and histological analysis showed unclear corticomedullary boundary **(Fig. 2k)**. The proportion of CD3^+^ cells in thymocytes of *Lig4^W447C/W447C^*mice were severely decreased compared with those of *Lig4^+/+^* mice (1.7% vs. 19%) **(Fig. 2l)**. Furthermore, there were very few T lineage cells, such as double-negative (DN) CD4^−^CD8^−^, double-positive (DP) CD4^+^CD8^+^, and single-positive (SP) CD4^+^ or CD8^+^ thymocytes **(Fig. 2m)**. The percentages of DN thymocytes in the *Lig4^W447C/W447C^* mice were increased, whereas those of DP and SP CD4^+^ thymocytes were decreased **(Fig. 2l and Extended Data Fig. 4a)**. In the spleen, CD3^+^CD4^+^ and CD3^+^CD8^+^ T cells were severely decreased in *Lig4^W447C/W447C^* mice **(Fig. 2g, n-p)**. Notably, CD4/CD8 ratio was higher in *Lig4^W447C/W447C^* mice than in *Lig4^+/+^* mice (**Fig. 2q**). The majority of CD4^+^ and CD8^+^ T cells remaining in the mutant mice were effector memory T cells (CD44^high^CD62L^low^) and naive T cells (CD44^low^CD62^high^) were hardly detected **(Fig. 2r, s)**. Concerning splenic Foxp3^+^ regulatory T cells (Tregs) in CD4^+^ T cells, the percentages were lower in *Lig4^W447C/W447C^* than in *Lig4^+/+^* mice **(Extended Data Fig. 4b, c)**. Taken together, *Lig4^W447C/W447C^* mice showed severe defects in both B and T cell generation, i.e. severe adaptive immunodeficiency.

### *Lig4^W447C/W447C^* mice develop severe intestinal inflammation

After weaning, *Lig4^W447C/W447C^* mice exhibited varying degrees of wasting, diarrhea, and rectal prolapse and showed higher mortality than *Lig4^+/+^* and *Lig4^W447C/+^* mice when housed in specific pathogen-free conditions. Kaplan-Meier analysis revealed that *Lig4^W447C/W447C^* mice had a median overall survival of 92 days **(Fig. 3a)**. In the macro-anatomical study, marked edema and shortening of the large intestine were found in almost all *Lig4^W447C/W447C^* mice around 10 weeks of age **(Fig. 3b)**. Histological examination of the large intestines revealed epithelial hyperplasia, goblet cell disappearance, and infiltration of inflammatory cells in the lamina propria and submucosa in *Lig4^W447C/W447C^*mice **(Fig. 3c).** Mouse colitis histology index (MCHI) of the *Lig4^W447C/W447C^*mice was significantly higher than that of *Lig4^+/+^* mice **(Fig 3d)**. Moreover, crypt abscesses, which are formed by active inflammation and are a hallmark of ulcerative colitis, were often present in the *Lig4^W447C/W447C^*mice **(Fig. 3c, arrowheads)**. Immunohistochemical analysis of the colons further showed that infiltrated cells in *Lig4^W447C/W447C^* mice consisted mainly of T cells, especially CD4^+^ T cells, and F4/80^+^ macrophages **(Fig. 3e).** These pathological findings were also observed in the small intestines of *Lig4^W447C/W447C^*mice **(Extended Data Fig. 5a, b)**. In *Lig4^W447C/W447C^*mice, increased T cells were mainly TCRαβ^+^ cells rather than TCRγδ^+^ cells, although their ratio of CD4 to CD8 was similar to that of T cells in *Lig4^+/+^* mice **(Extended Data Fig. 5c, d).** Meanwhile, B220^+^ cells were hardly detected (**Fig. 3e and Extended Data Fig. 5a**). *Lig4^W447C/W447C^* mice therefore showed severe inflammations in the small and large intestines with infiltration mainly of CD4+ T cells and macrophages into the mucosa and submucosa. We also analyzed the gut microbiota by fecal 16S rRNA gene sequencing. The composition of the gut microbiota was comparable between *Lig4^+/+^* and *Lig4^W447C/W447C^*mice **(Extended Data Fig. 6)**. The results indicate that the cause of intestinal inflammations is unlikely due to the expansion of certain bacteria.

**Figure 3.**
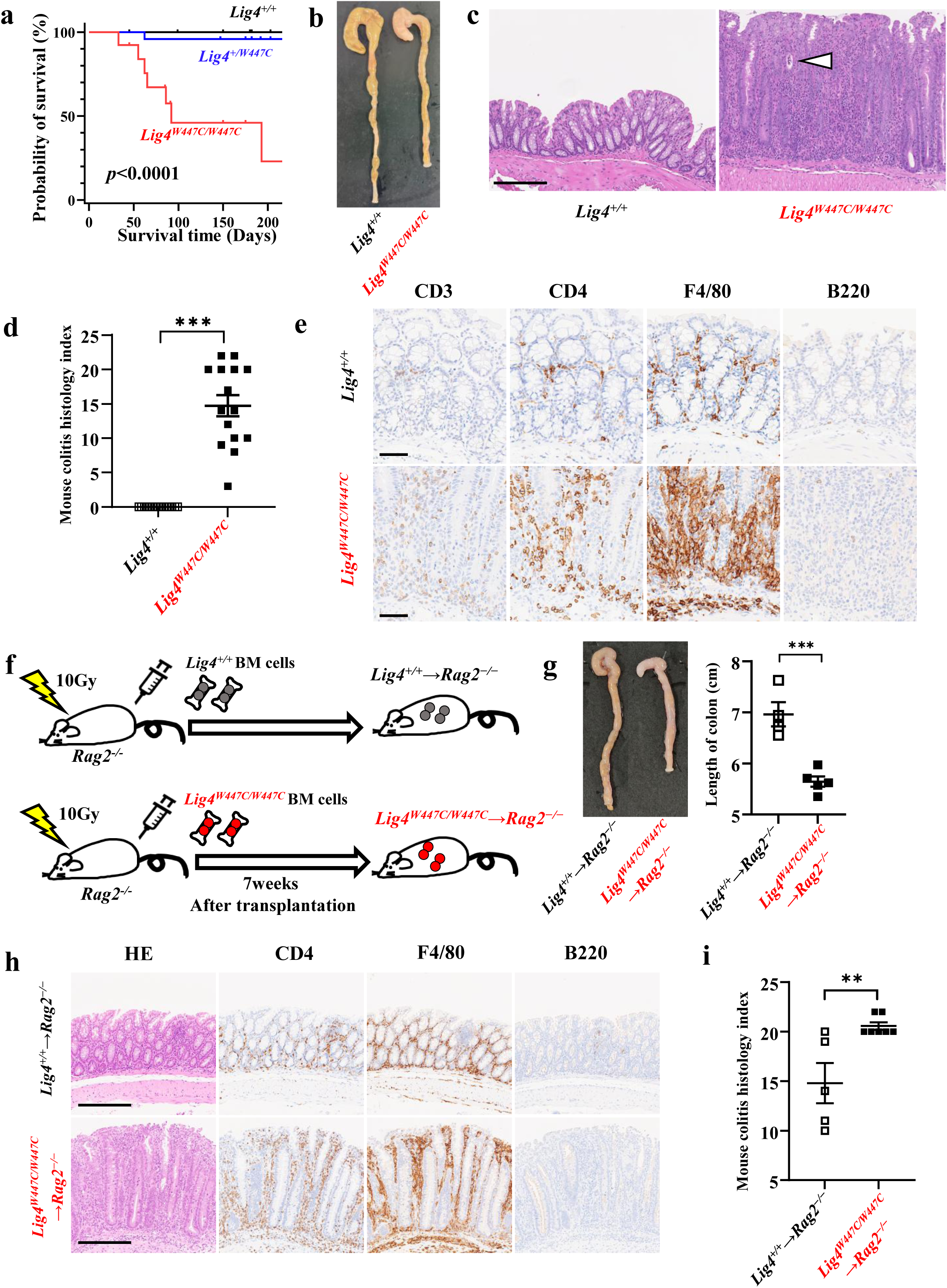
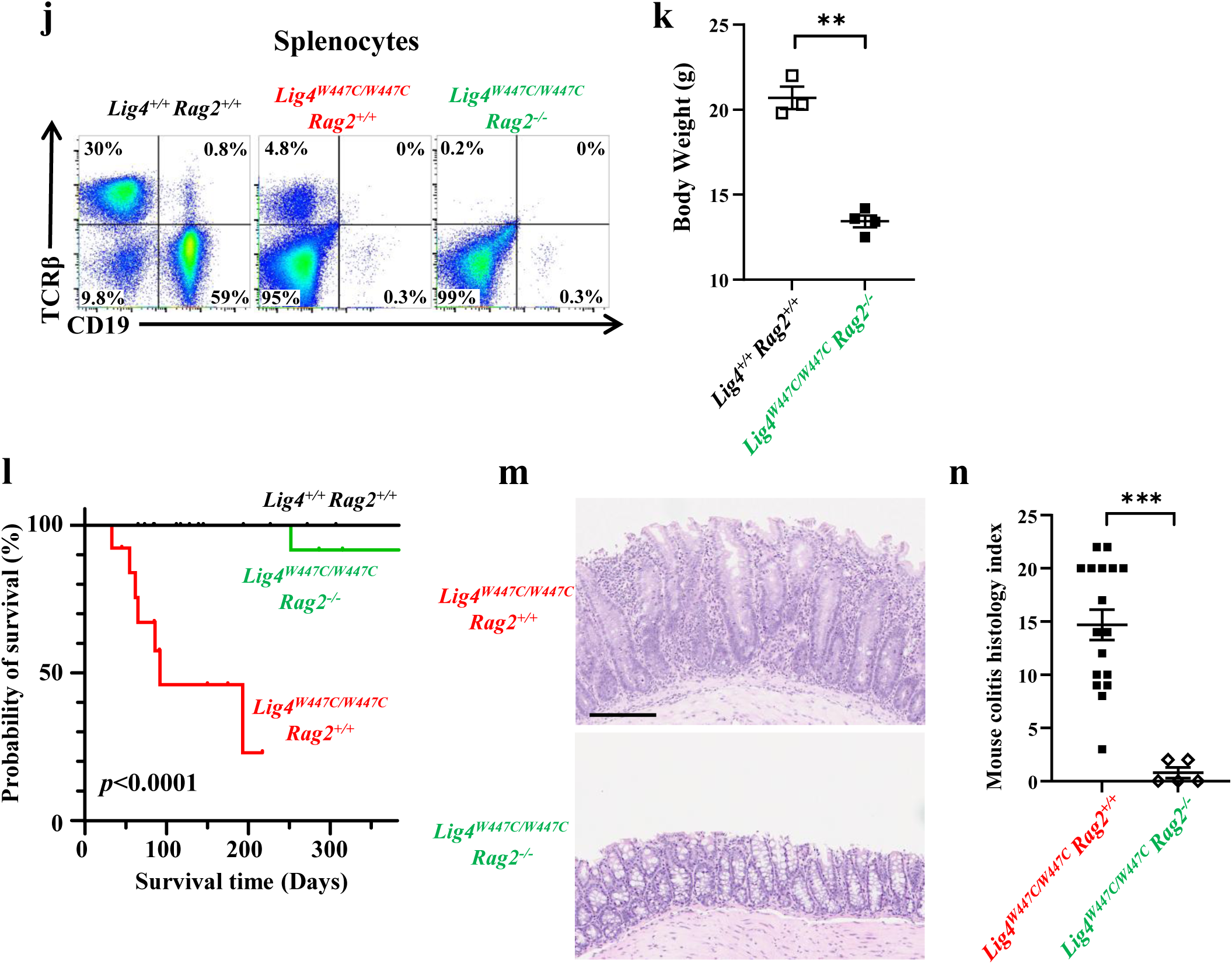
Hematopoietic cells caused severe intestinal inflammation in *Lig4^W447C/W447C^* mice. (a) Kaplan–Meier survival curves of *Lig4^+/+^* (*n* = 13), *Lig4^+/W447C^* (*n* = 23), and *Lig4^W447C/W447C^*(*n* = 12) mice. Statistical analysis was performed using the log-rank test. (b) Macroscopic features of the *Lig4^W447C/W447C^* mouse colon revealed an edematous, thickened, and shortened colon. (c) Representative H&E staining of colons from *Lig4^+/+^* and *Lig4^W447C/W447C^* mice. White arrow indicates a crypt abscess. Scale bar represents 200 μm. (d) Mouse colitis histology index of *Lig4^+/+^* and *Lig4^W447C/W447C^* mice (*Lig4^+/+^* [*n* = 15] and *Lig4^W447C/W447C^* [*n* = 15]; 7–13 weeks). (e) Immunohistochemistry of colons from *Lig4^+/+^* and *Lig4^W447C/W447C^* mice with CD3, CD4, F4/80 and B220 antibodies. Scale bar represents 60 μm. (f) The scheme of BM transplantation experiment. BM cells from *Lig4^+/+^* or *Lig4^W447C/W447C^* mice were transplanted into irradiated *Rag2^−/−^* mice. (g) A shortened and edematous colon from a *Lig4^+/+^*→*Rag2*^−/−^ mouse and a *Lig4^W447C/W447C^*→*Rag2^−/−^* mouse. The graph (right) depicts the length of colons. (*Lig4^+/+^*→*Rag2^−/−^* [*n* = 4] and *Lig4^W447C/W447C^*→*Rag2^−/−^* [*n* = 5]). (h) H&E staining and immunohistochemistry with CD4, F4/80 and B220 antibodies of colons from a *Lig4^+/+^*→*Rag2^−/−^* mouse and a *Lig4^W447C/W447C^*→*Rag2^−/−^* mouse. Scale bar represents 100 μm. (i) Mouse colitis histology index of *Lig4^+/+^*→*Rag2^−/−^* and *Lig4^W447C/W447C^*→*Rag2^−/−^* mice. (*Lig4^+/+^*→*Rag2^−/−^* [*n* = 5] and *Lig4^W447C/W447C^*→*Rag2^−/−^* [*n* = 7]). (j) FACS analysis of CD19 and TCRβ expression in splenocytes from *Lig4^+/+^Rag2^+/+^*, *Lig4^W447C/W447C^Rag2***^+/+^** and *Lig4^W447C/W447C^Rag2**^−/−^***mice. (k) The graph depicts the weight of female mice (*Lig4^+/+^Rag2^+/+^* [*n* = 3] and *Lig4*^W447C/W447C^*Rag2^−/−^* [*n* = 4]; 14–16 weeks). (l) Kaplan–Meier survival curves of *Lig4^+/+^Rag2^+/+^*(*n* =21), *Lig4^W447C/W447C^Rag2^+/+^* (*n* =13), and *Lig4^W447C/W447C^Rag2**^−/−^*** (*n* =12) mice. Statistical analysis was performed using the log-rank test. (m) Representative H&E staining of colons from *Lig4^W447C/W447C^Rag2^+/+^* and *Lig4^W447C/W447C^Rag2**^−/−^*** mice. Scale bar represents 200 μm. (n) Mouse colitis histology index of *Lig4^W447C/W447C^Rag2^+/+^* and *Lig4^W447C/W447C^Rag2^−/−^* mice (*Lig4^W447C/W447C^Rag2^+/+^* [*n* = 17] and *Lig4^W447C/W447C^Rag2^−/−^*[*n* = 5]; 7– 18 weeks). The data are presented as the mean ± standard error of the mean (SEM).

### Intestinal inflammation in *Lig4^W447C/W447C^* mice is dependent on lymphocytes

Next, to determine whether hematopoietic cells were sufficient for the development of colitis in the *Lig4^W447C/W447C^* mice, we generated BM chimeric mice, by transferring BM cells from *Lig4^+/+^* and *Lig4^W447C/W447C^*mice into irradiated *Rag2*^−/−^ mice, which are hereafter referred as *Lig4^+/+^*→*Rag2*^−/−^ and *Lig4^W447C/W447C^*→*Rag2*^−/−^mice, respectively **(Fig. 3f)**. Compared with *Lig4^+/+^*→*Rag2*^−/−^mice, *Lig4*^W447C/W447C^→*Rag2*^−/−^mice showed severe intestinal manifestations, including diarrhea, colonic shortness, and edema **(Fig. 3g)**. Furthermore, in the colons of *Lig4^W447C/W447C^*→*Rag2^−/−^* mice, CD4^+^ T cells and macrophages mainly infiltrated as observed in those of *Lig4^W447C/W447C^* mice **(Fig. 3h, i)**.

These results suggest that hematopoietic cells are responsible for the development of intestinal inflammations in *Lig4^W447C/W447C^*mice. Meanwhile, *Lig4^W447C/W447C^* mice were sensitive to and succumbed to irradiation. Therefore, we could not generate BM chimeric mice by using *Lig4^W447C/W447C^* mice as recipients. Thus, we could not assess the roles of non-hematopoietic cells in intestinal inflammations in *Lig4^W447C/W447C^* mice.

To further investigate the involvement of lymphocytes in intestinal inflammations in the *Lig4^W447C/W447C^* mice, *Lig4^W447C/W447C^Rag2^−/−^*mice were generated by crossing *Lig4^W447C/W447C^* mice with *Rag2^−/−^* mice. Splenic T cells were still detected in *Lig4^W447C/W447C^*mice, but they were absent in *Lig4^W447C/W447C^Rag2^−/−^*mice **(Fig. 3j)**. The double mutant mice had growth disturbances and microcephaly but showed significantly longer survival than *Lig4^W447C/W447C^* mice **(Fig. 3k, l)**. Notably, *Lig4^W447C/W447C^Rag2*^−/−^ mice developed no signs of intestinal inflammations **(Fig. 3m, n)**. These results suggest that lymphocytes are required for the development of intestinal inflammations in *Lig4^W447C/W447C^*mice.

### Expression of IFN-γ inducible genes were enhanced in the colon of *Lig4^W447C/W447C^*mice

To characterize intestinal inflammations in *Lig4^W447C/W447C^* mice, RNA-sequence (RNA-Seq) analysis was performed on colonic tissues from *Lig4^+/+^* and *Lig4^W447C/W447C^* mice. In total, 25,706 genes were differentially expressed between *Lig4^+/+^* and *Lig4^W447C/W447C^* mice (**Supplementary Table 1**). Expression of 2278 and 3728 genes was up– and down-regulated, respectively, more than 1.5-fold. Unbiased gene set enrichment analysis (GSEA) of these genes revealed up– and down-regulation of 24 and two gene sets, respectively (nominal *p*-value < 0.05, false discovery rate q-value < 0.25) **(Fig. 4b and Supplementary Table 2)**. The most upregulated gene set was ‘interferon gamma response’ (normalized enrichment score=2.97; *p*-value < 0.001 **(Fig. 4a-c)**. Expression of 15 representative IFN-γ-inducible genes such as *Ciita* or *Cxcl9* was more than five-fold higher in the colons of *Lig4^W447C/W447C^* mice than in those of *Lig4^+/+^* mice (**Fig. 4a and Supplementary Table 1**).

**Figure 4.**
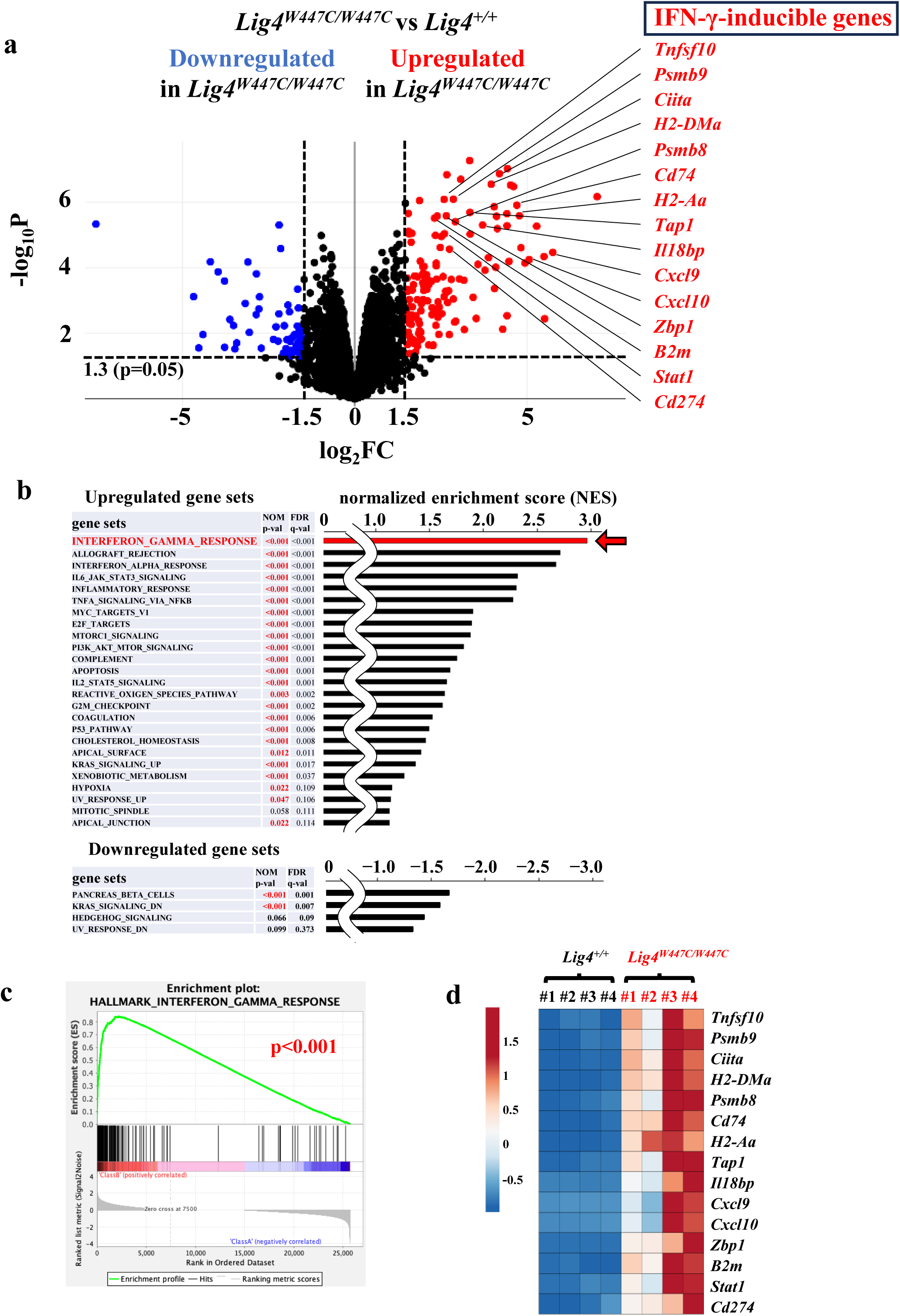
Bulk RNA-sequencing analysis of the colon of *Lig4^W447C/W447C^*mice. (a) Relative expression levels of the colon from eight 10-week-old mice (*Lig4^+/+^* [*n* = 4] and *Lig4^W447C/W447C^*[*n* = 4]) analyzed by bulk RNA-sequencing (RNA-seq) analysis. Visualization of bulk RNA-seq results with a volcano plot. The dotted lines indicate the value of fold changes (±^21.5^-fold, x-axis) and the significance p-value (*p*<0.05, y-axis). (b) Normalized enrichment scores of excerpted top 25 up-regulated (upper) and top 4 down-regulated (bottom) gene sets in the colons from *Lig4^W447C/W447C^* mice compared with the *Lig4^+/+^* mice analyzed by gene set enrichment analysis (GSEA). Gene sets associated interferon-gamma response, indicated by the red arrow, represent the focus of this study. “NOM p-val” represents the nominal p-value, and “FDR q-val” represents the false discovery rate q-value. (c) GSEA enrichment plot for “interferon-gamma response” with comparison of the colons from four *Lig4^W447C/W447C^*mice and four *Lig4^+/+^* mice. “*p*” indicates nominal *p*-value. (d) The heatmap shows the changes in gene expression in the IFN-γ-inducible genes within the “interferon-gamma response” gene sets. Color ranges from dark red to dark blue representing the highest and lowest expression of a gene, respectively.

Furthermore, expression of these genes was more prominently enhanced in the colons from *Lig4^W447C/W447C^* mice with severe manifestations (MCHI: #3-score 17, #4-score 20) than those with relatively mild manifestations (MCHI: #1-score 9, #2-score 3) (**Fig. 4d**). Thus, intestinal inflammations in *Lig4^W447C/W447C^* mice are featured by enhanced production of IFN-γ.

### Loss of IFN-**γ** expression in *Lig4^W447C/W447C^*mice evoked Th2/Th17-mediated intestinal inflammation

IFN-γ is a key cytokine expressed in Th1 cells among lymphocytes. We therefore compared the expression of various Th cell subset signature genes. Th1 cell signature genes including *Ifng* were expressed at significantly higher levels in *Lig4^W447C/W447C^*colons than in *Lig4^+/+^* colons **(Fig. 5a)**. Expression of Th2 and Treg cell signature genes were also increased, whereas the expression of some Th17 cell signature genes was rather decreased in the mutant colons **(Fig. 5a)**.

**Figure 5.**
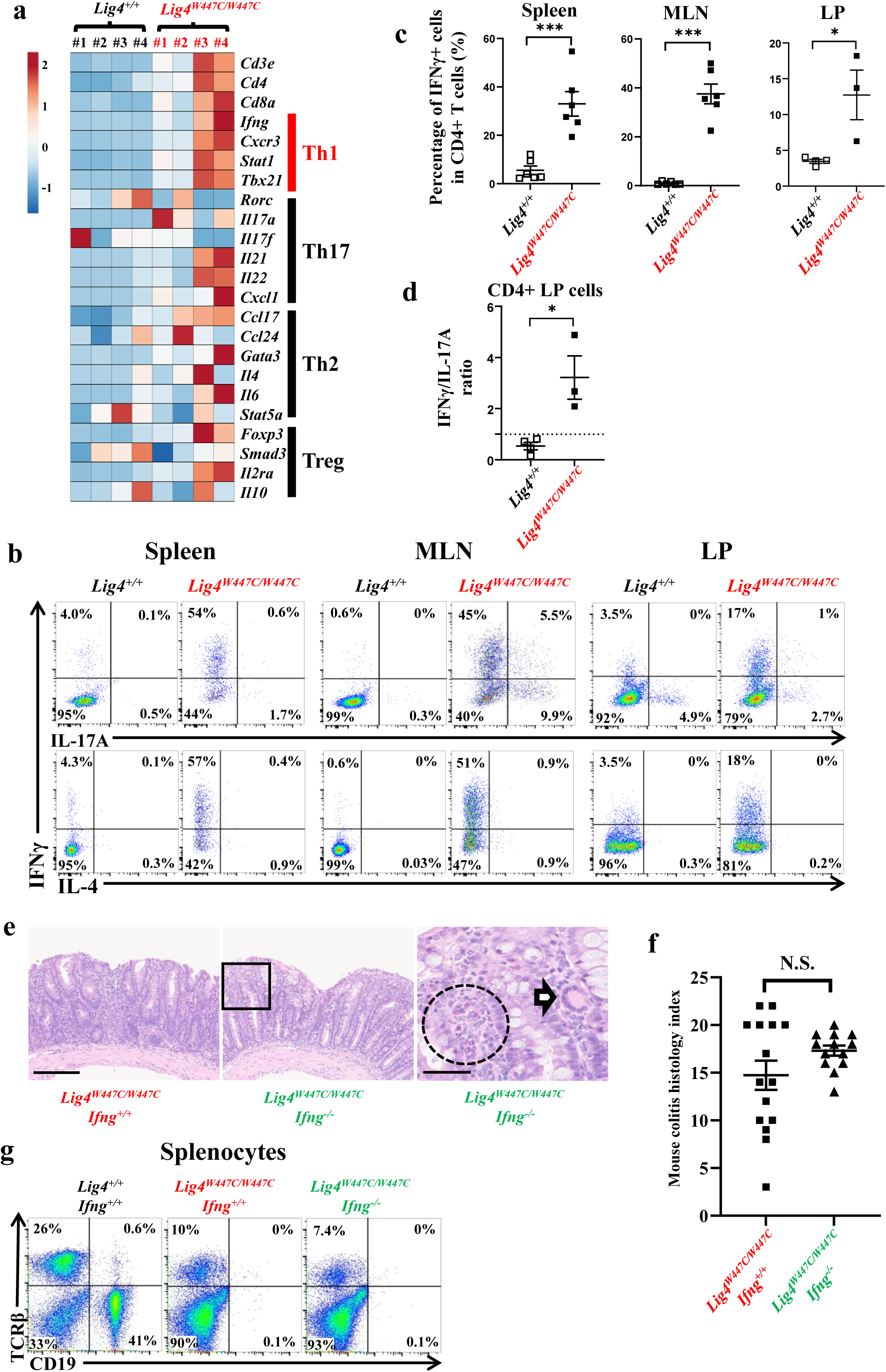
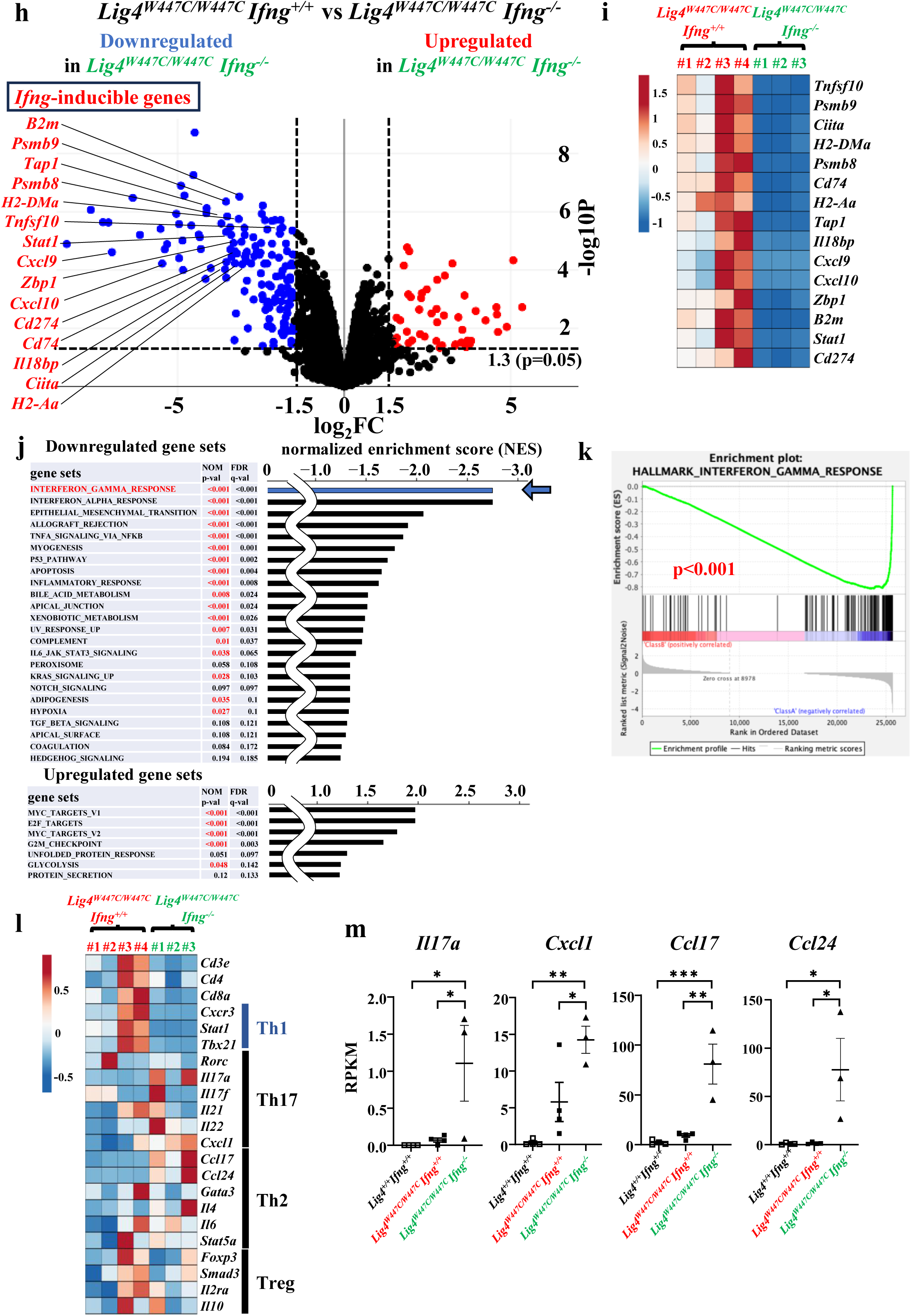
Analysis of *Lig4^W447C/W447C^Ifng^−/−^* mice. (a) The heatmap shows the changes in gene expression of Th cell subset signature genes in the colon from four 10-week-old mice (*Lig4^+/+^* [*n* = 4] and *Lig4^W447C/W447C^*[*n* = 4]) analyzed by bulk RNA-sequencing (RNA-seq) analysis. Color ranges from dark red to dark blue representing the highest and lowest expression of a gene, respectively. (b) Intracellular cytokine staining (IFN-γ, interleukin (IL)-17A, and IL-4) of CD4^+^ T cells in splenocytes, mesenteric lymph nodes (MLN), and intestinal lamina propria (LP) from *Lig4^+/+^* and *Lig4^W447C/W447C^* mice. Samples were stimulated with PMA (50 ng/mL), ionomycin (500 ng/mL) and Golgi plug for 5 hours. (*Lig4^+/+^* [Spleen and MLN: *n* = 6, LP: *n* = 4] and *Lig4^W447C/W447C^*[Spleen and MLN: *n* = 6, LP: *n* = 3]). (c) The percentages of IFN-γ-producing CD4^+^ T cells in the respective tissues. (d) The ratio of IFN-γ/IL-17A-producing CD4^+^ T cells in LP from each mouse. (e) Representative H&E staining of colons from *Lig4^W447C/W447C^Ifng^+/+^*(left panel) and *Lig4^W447C/W447C^Ifng^−/−^* mice (middle and right panels). In middle panel, black square line denotes the areas that are presented magnified in the right panel. In the magnified image, infiltrated eosinophils (dotted circle line) and multinucleated giant cell (arrow) were found in *Lig4^W447C/W447C^Ifng^−/−^* mice. Scale bar represents 200 μm (left and middle panels) and 50 μm (right panel). (f) The graph depicts the mouse colitis histology index of the representative mice (*Lig4^W447C/W447C^Ifng^+/+^* [*n* = 15], and *Lig4^W447C/W447C^Ifng^−/−^* [*n* = 13]; 7-13 weeks). (g) FACS analysis of CD19 and TCRβ expression in splenocytes from *Lig4^+/+^Ifng^+/+^*, *Lig4^W447C/W447C^Ifng^+/+^* and *Lig4^W447C/W447C^Ifng^−/−^*mice. (h) Relative expression levels of the colon from seven 10-week-old mice (*Lig4^W447C/W447C^Ifng^+/+^*[*n* = 4] and *Lig4^W447C/W447C^Ifng^−/−^*[*n* = 3]) analyzed by bulk RNA-seq analysis. Visualization of bulk RNA-seq results with a volcano plot. The dotted lines indicate the value of fold changes (±^21.5^-fold, x-axis) and the significance *p*-value (*p*<0.05, y-axis). (i) The heatmap shows the changes in gene expression in the IFN-γ-inducible genes within the ‘interferon-gamma response’ gene sets among colons from *Lig4^W447C/W447C^Ifng^+/+^* and *Lig4^W447C/W447C^Ifng^−/−^* mice. Color ranges from dark red to dark blue representing the highest and lowest expression of a gene, respectively. (j) Normalized enrichment scores of excerpted tops 24 down-regulated (upper) and seven up-regulated (bottom) gene sets in the colons from the *Lig4^W447C/W447C^Ifng^−/−^* mice [*n* = 3] compared with *Lig4^W447C/W447C^Ifng^+/+^* mice [*n* = 4] analyzed by gene set enrichment analysis (GSEA). Gene sets associated ‘interferon-gamma response’, indicated by the blue arrow, represent the focus of this study. ‘NOM *p*-val’ represents the nominal p-value, and ‘FDR q-val’ represents the false discovery rate q-value. (k) GSEA enrichment plot for ‘interferon-gamma response’ with comparison of the colons from four *Lig4^W447C/W447C^Ifng^+/+^*mice and three *Lig4^W447C/W447C^Ifng^−/−^* mice. ‘*p*’ indicates nominal *p*-value. (l) The heatmap shows the changes in representative gene expressions of Th cell subset signature genes among colons from *Lig4^W447C/W447C^Ifng^+/+^*and *Lig4^W447C/W447C^Ifng^−/−^* mice. (m) The reads per kilobase per million (RPKM) levels of the *Il17a, Cxcl1, Ccl17 and Ccl24* genes in the colon from *Lig4^+/+^Ifng^+/+^*, *Lig4^W447C/W447C^Ifng^+/+^*and *Lig4^W447C/W447C^Ifng^−/−^* mice analyzed by bulk RNA-seq analysis. The data are presented as the mean ± standard error of the mean (SEM).

We have also performed intracellular cytokine staining analysis in CD4^+^ T cells from *Lig4^+/+^* and *Lig4^W447C/W447C^*mice. IFN-γ-expressing CD4^+^ T cells was higher in frequency in the spleen, mesenteric lymph nodes (MLNs), and lamina propria (LP) of *Lig4^W447C/W447C^* mice than in those of *Lig4^+/+^*mice **(Fig. 5b, c)**. Notably, almost half of CD4^+^ T cells expressed IFN-γ in *Lig4^W447C/W447C^*mice. Meanwhile, IL-4-expressing CD4^+^ T cells were comparable **(Fig. 5b)**. IL-17A-expressing CD4^+^ T cells were increased in the mutant MLN and LP lymphocytes than in the *Lig4^+/+^*lymphocytes, but their proportions were less than 10% **(Fig. 5b)**. Among LP CD4^+^ T cells, IL-17A-expressing cells were dominant in *Lig4^+/+^* mice, whereas IFN-γ-expressing cells were dominant in the mutant mice **(Fig. 5d)**. Dominance of IFN-γ-expressing cells were more prominent in the spleen and MLN CD4^+^ T cells in *Lig4^W447C/W447C^*mice. These results suggest that Th1 cells producing IFN-γ are mainly activated in *Lig4^W447C/W447C^* mice.

To determine whether and how IFN-γ is involved in the development of intestinal inflammations in *Lig4^W447C/W447C^* mice, we generated *Lig4^W447C/W447C^Ifng^−/−^* mice. The double mutant mice showed developmental defect and microcephaly, in similar to *Lig4^W447C/W447C^* mice. The double mutant mice had inflammatory cell infiltration in the intestine, with a similar inflammatory score to that of the *Lig4^W447C/W447C^* mice (**Fig. 5e, f**). These inflammatory cells were mainly neutrophils in the LP and crypt abscesses were not found **(Fig. 5e)**. Furthermore, eosinophils were often found in the lesions (**Fig. 5e, dotted circle**) and multinucleated giant cells, which could be potentially derived from activated macrophages, were occasionally found (**Fig. 5e, arrow**). Flow cytometric analysis revealed residual T cells in the splenocytes of *Lig4^W447C/W447C^Ifng^−/−^*mice (**Fig. 5g**). Intracellular staining analysis of the residual T cells lacked expression of IFN-γ, but increased expression of IL-4 and IL-17A (**Extended Data Fig. 7**). The transcriptome analysis also revealed that expression of IFN-γ inducible genes was profoundly decreased in the colons of *Lig4^W447C/W447C^Ifng^−/−^*mice compared with *Lig4^W447C/W447C^* mice (**Fig. 5h, i and Supplementary Table 1**). GSEA comparing *Lig4^W447C/W447C^Ifng^−/−^* and *Lig4^W447C/W447C^* mice showed upregulation and downregulation of 18 and 5 gene sets (nominal p-value < 0.05, false discovery rate q-value < 0.25), respectively, and confirmed the most downregulated gene set was ‘interferon-gamma response’ (**Fig. 5j, k and Supplementary Table 2**). Furthermore, we compared the expression of various Th cell subset signature genes. Expression of Th1 cell signature genes in the *Lig4^W447C/W447C^Ifng^−/−^*colons was returned to comparable levels to that in the *Lig4^+/+^Ifng^+/+^*colons **(Supplementary Table 1)**. Meanwhile, expression of Th17 (*Il17a, Cxcl1*) and Th2 (*Ccl17* and *Cc124*) cell signature genes were upregulated in *Lig4^W447C/W447C^Ifng^−/−^* colons, compared with *Lig4^+/+^Ifng^+/+^* and *Lig4^W447C/W447C^Ifng^−/−^* colons (**Fig. 5l, m and Supplementary Table 1**). Expression of Treg signature genes was comparable among these mice (**Fig. 5l and Supplementary Table 1**). Taken together, IFN-γ deficiency ameliorated Th1-skewed intestinal inflammations, but instead deteriorated Th17-skewed intestinal inflammations in *Lig4^W447C/W447C^* mice.

### *Lig4^W447C/W447C^* mice exhibit a biased TCR repertoire in the spleen

We then performed a comprehensive bulk-based analysis of a diverse repertoire of T cell receptors in the spleen and the MLN to investigate whether T cell repertoire shows some skewing in *Lig4^W447C/W447C^* mice. The T cell repertoire was quite heterogeneous in *Lig4^+/+^* mice, while certain T cell receptors were abundantly expressed in *Lig4^W447C/W447C^* mice **(Extended Data Fig. 8a-d and Supplementary Table 3, 4)**. A close examination of the 3’ (proximal) Vα segments revealed a striking difference in their usages among *Lig4^+/+^* and *Lig4^W447C/W447C^* mice **(Extended Data Fig. 8a, b)**. TCR Vα usage in the spleen and the MLN of *Lig4^W447C/W447C^* mice was strongly skewed to the 3’ (proximal) end. Meanwhile, as for Vβ, a slight bias in the utilization frequency of segments was observed in *Lig4^W447C/W447C^* mice, regardless of their position (**Extended Data Fig. 8c, d**).

Next, we performed single-cell RNA-Seq (scRNA-seq) analysis to investigate which pairs of T cell receptors are expressed in the spleens of *Lig4^+/+^* and *Lig4^W447C/W447C^*mice. Consistent with flow cytometry analysis, T cells were severely decreased, and the residual T cells mainly consisted of Th1 cells in the spleens of the mutant mice **(Fig. 6a, b)**. *Lig4^+/+^* T cells from 2015 barcodes included 2002 clonotypes, which contained the clones limited to two at most, and all but the top six clonotypes were represented by one cell. Meanwhile, 667 *Lig4^W447C/W447C^*T cells consisted of 235 clonotypes and the top 10 clonotypes each occupied more than 10 T cells, 1.5% of the total T cells **(experiment [exp] #1 in Fig. 6c)**. Those 10 clones contained four CD4^+^ cells, five CD8^+^ cells, and one CD4^−^CD8^−^ T cell. Seven out of 10 clones did not express *Sell*, indicating that most of them are activated. Two CD4^+^ T cell clonotypes, which are activated and express *Trbv3/Trbj1-1* or *Trbv16/Trbj2-3*, showed high expression of *Ifng* **(exp #1 in Fig. 6d, e**). Meanwhile, neither of these 10 T cell clonotypes expressed *Il4* or *Il17a* **(exp #1 in Fig. 6d)**. Furthermore, we analyzed the T cell repertoire diversity in additional two *Lig4^W447C/W447C^* spleens (exp #2 and #3) by scRNA-seq analysis. In exp #2 and #3 mutant mice, 1,265 and 1,610 T cells consisted of 366 and 416 clonotypes, respectively (Fig. 6c). These top 10 clonotypes each occupied 1.26% and 1.30% of the total T cells **(exp #2 and #3 in Fig. 6c)**. In #2 mutant mouse, two CD4^+^ T cell clonotypes expressing *Ifng* represented *Trbv3/Trbj2-3/Trav14-3/Traj56* or *Trbv15/Trbj2-5//Trav13-1/Traj58* transcripts **(exp #2 in Fig. 6d, e)**. Meanwhile, #3 mutant mouse predicted all effector memory CD4^+^ T cell among top 9 clonotypes; however, none of them expressed *Ifng*, *Il4* or *Il17a* **(exp #3 in Fig. 6d, e)**. Therefore, several T cell clonotypes, which might include pathogenic ones, were skewed and expanded in *Lig4^W447C/W447C^*mice.

**Figure 6.**
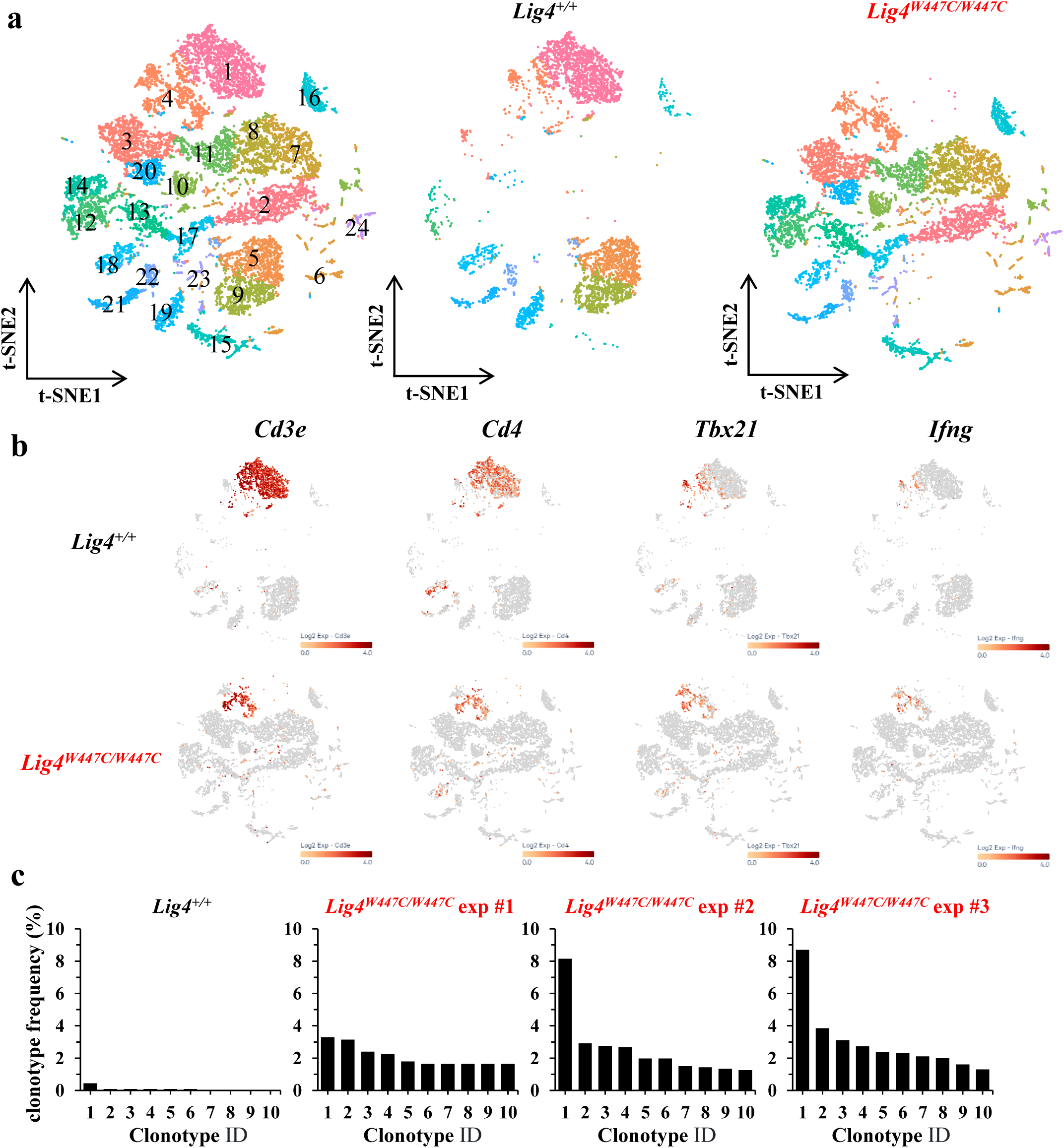
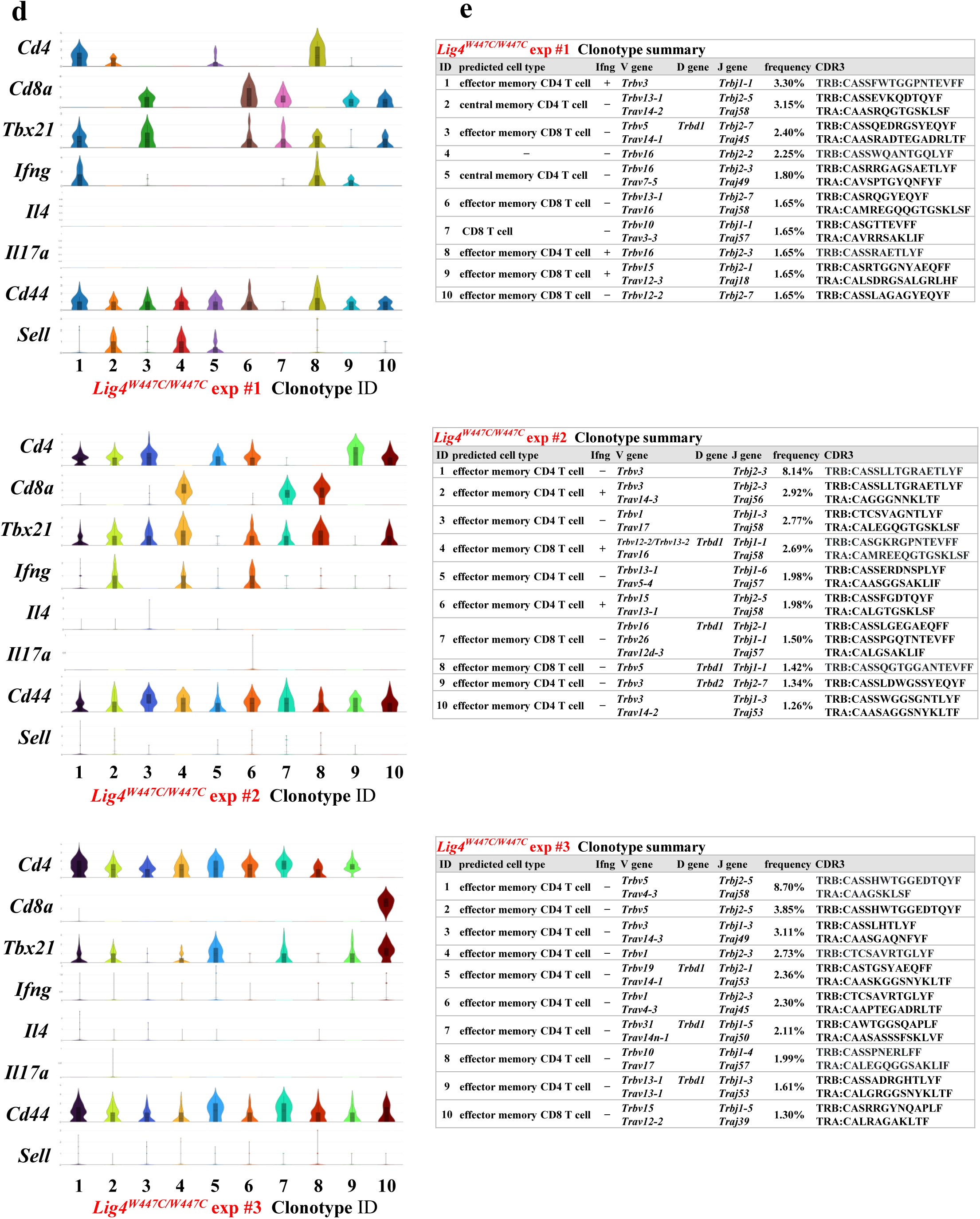
Single-cell RNA-sequencing of splenocytes samples. (a) The t-SNE plots of splenocytes from *Lig4^+/+^* and *Lig4^W447C/W447C^*mice based on 17,377 cells (*Lig4^+/+^*: 5,289 cells, *Lig4^W447C/W447C^*: 12,088 cells). It was classified into 24 clusters. t-SNE plots show the data of total (left), *Lig4^+/+^* (middle) and *Lig4^W447C/W447C^*(right). (b) Feature plots of *Cd3e*, *Cd4*, *Tbx21* and *Ifng*. The color bar, from yellow to brown, reveals gradual expression intensity differences from low to high. (c) The clonotype frequencies of splenocytes from *Lig4^+/+^* mice and *Lig4^W447C/W447C^*mice [n=3; exp #1, #2 and #3]. (d) The violin plots of the expression of the listed gene in the respective clonotype of each *Lig4^W447C/W447C^*mice [n=3; exp #1, #2 and #3]. (e) The clonotype summary of each *Lig4^W447C/W447C^* mice [n=3; exp #1, #2 and #3].

Moreover, we quantitated usage of TCR V and J gene segments (Vα, Jα, Vβ, and Jβ) in CDR3 sequences by scRNA-Seq analysis in splenic T cells of one *Lig4^+/+^*and three *Lig4^W447C/W447C^* mice **(Figure 7a-d and Extended Data** Fig 9a, b). Vα repertoire in T cells from three mutant mice was dominated by the most 3’ (proximal) segments from *Trav16* to *Trav21-dv12* segments (**Fig 7a and Extended Data Fig. 9a**). Conversely, the Jα repertoire in the mutant T cells were dominated by the most 5’ (distal) segments from *Traj58* to *Traj49* **(Fig 7b and Extended Data Fig. 9a)**. Moreover, Vβ and Jβ segments in the mutant T cells were evenly distributed, although usage of some segments, including *Trbv1*, *Trbv3*, and *Trbj1-1*. was increased **(Fig 7c, d and Extended Data Fig. 9b**).

**Figure 7.**
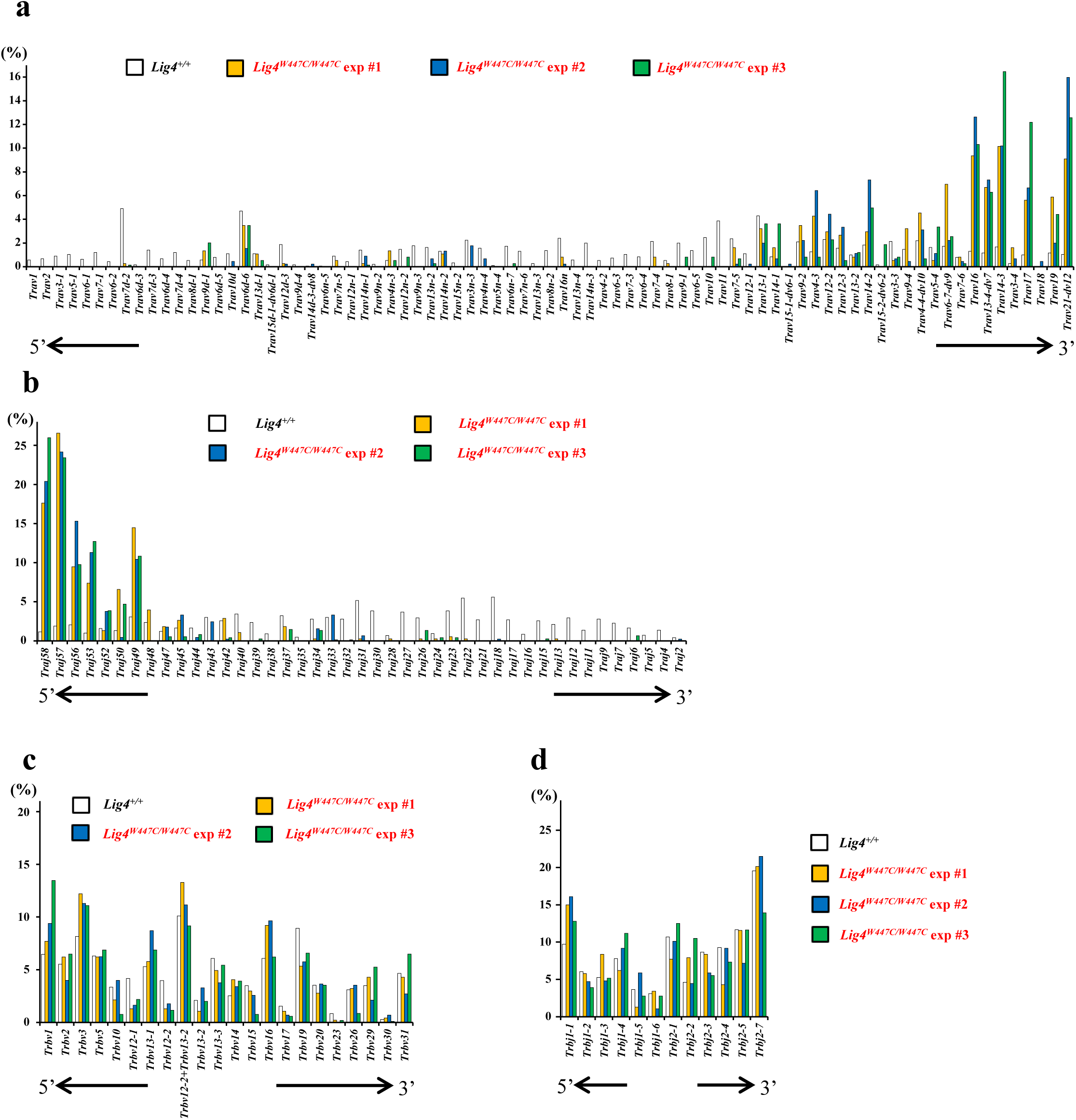
TCR repertoire analysis by single-cell RNA-sequencing of splenocytes samples. The bar graphs depict the utilization frequencies of TCR gene segments in *Lig4^+/+^* mice [n=1] and *Lig4^W447C/W447C^* mice [n=3; exp #1, #2 and #3]. (a) *Trav* segments, arranged in chromosomal order from the 5’ end (left) to the 3’ end (right) of the *Trav* locus. (b) *Traj* segments, arranged in chromosomal order from the 5’ end (left) to the 3’ end (right) of the *Traj* locus. (c) *Trbv* segments, arranged in chromosomal order from the 5’ end (left) to the 3’ end (right) of the *Trbv* locus. (d) *Trbj* segments, arranged in chromosomal order from the 5’ end (left) to the 3’ end (right) of the *Trbj* locus.

## Discussion

We introduced a novel hypomorphic gene variant, p.W447C, derived from a patient with LIG4 syndrome into mice. The homozygous mutant mice showed not only developmental and neuronal defects but also severe adaptive immunodeficiency. They further showed severe intestinal inflammations with intestinal epithelial hyperplasia, marked infiltration of Th cells and macrophages, and cryptic abscesses, which are characteristic of ulcerative colitis. The inflammations were developed in *Lig4^W447C/W447C^* →*Rag2^−/−^* mice and abolished in *Lig4^W447C/W447C^ Rag2^−/−^* mice. Furthermore, the lesions showed high expression of IFN-γ and IFN-γ-induced genes and were skewed towards Th2/Th17-type inflammations by IFN-γ deficiency. These results indicate that intestinal inflammations in *Lig4^W447C/W447C^*mice are driven by Th1 cells.

LIGIV-deficient mice are embryonic lethal. Meanwhile, so far two kinds of hypomorphic *Lig4* homozygous mutant mice, Y288C and R278H, were generated and found to be born with growth retardation^16, 17, 18, 19, 20, 21^. Both *Lig4^Y288C/Y288C^*and *Lig4^R278H/R278H^* mice manifested defective DSB repair and adaptive immunodeficiency, as observed also in *Lig4^W447C/W447C^*mice ^17, 19, 20^. However, the immunodeficiency in these two mutant mice was milder than that in *Lig4^W447C/W447C^*mice. Although protein expression of LIGIV is decreased in *Lig4^Y288C/Y288C^*and *Lig4^R278H/R278H^* mice, it is preserved in *Lig4^W447C/W447C^*mice ^19, 21^. This indicates that protein stability or synthesis is impaired by Y288C and R278H, but not by W447C mutations, and it indicates that defective protein expression caused by Y288C and R278H mutations should contribute to functional defects of LIGIV. It remains unknown why W447C mutation, which does not reduce LIGIV expression, caused more severe adaptive immunodeficiency than the two other mutations. Y288, R278 and W447 are all localized in the middle of the enzymatic domain. According to *in silico* analysis, neither Y288C, R278H, or W447C mutations disturb the overall structure of LIGIV. In order to repair DSB, LIGIV should first bind to AMP ^23^. W447 faces to AMP and W447C mutation is supposed to cause decrease of the interface area and enlargement of the AMP-binding pocket, which should then lead to destabilization of the interaction of LIGIV and AMP. Although R278 also faces to AMP and R278H mutation leads to decrease of the interface area, the mutation minimally affects the AMP-binding pocket. Y288 is located on the surface of LIGIV and is not directly involved in the interaction of LIGIV with AMP. The AMP-binding pocket should therefore be structurally kept by Y288C mutation. Based on these findings, it can be assumed that W447C mutation should disturb the interaction of LIGIV with AMP more severely than the other two mutations, thereby leading to more severe defects in adaptive immunity.

Among three hypomorphic *Lig4* homozygous mutants, *Lig4^W447C/W447C^*mice are featured by their inflammatory phenotype. Meanwhile, *Lig4^R278H/R278H^*and *Lig4^Y288C/Y288C^* mice showed increased morbidity and high incidence of thymic tumors, indicating that these two mutations led to increased susceptibility to malignancy ^19, 20^. In this study, we could not formally assess whether *Lig4^W447C/W447C^* mice manifest the lymphoid malignancies, because most of them do not survive long enough to be analyzed. No apparent inflammatory conditions have been found in these previously reported mice, although only a modest infiltration of T lymphocytes and neutrophils was observed in the guts and livers of *Lig4^R278H/R278H^*mice ^17, 19^. Furthermore, splenic T cells in the *Lig4^R278H/R278H^*mice did not show any prominent skewing in their cytokine expression profiles ^19^. Thus, Th1 cell-driven intestinal inflammation with macrophages in the *Lig4^W447C/W447C^* mice is a unique phenotype.

The RAG proteins, involved in V(D)J recombination of BCRs and TCRs, are essential for lymphocyte development. Several hypomorphic mutations in *Rag* genes also lead to inflammatory manifestations driven by lymphocytes with adaptive immunodeficiency, which are called as Omenn syndrome ^24, 25, 26, 27, 28^. Among several *Rag* mutant mice, *Rag2^R229Q/R229Q^* mice have manifested inflammatory conditions, including colitis, which are similar to intestinal inflammations in *Lig4^W447C/W447C^* mice in terms of dominant infiltration of Th cells ^29^. However, *Rag2^R229Q/R229Q^* mice show not only Th1-, but also Th17– and Th2-driven inflammations, which are characterized by skin lesions with erythroderma and eosinophilia ^29, 30, 31^. Meanwhile, in *Lig4^W447C/W447C^* mice, the intestinal inflammations were driven mainly by Th1 cells and skin lesions were not observed. Furthermore, B lineage cells were hardly detected in the intestines or spleen and serum Ig levels including IgE and anti-dsDNA Abs were low, while levels of serum IgM and IgGs were kept and the levels of serum IgE and anti-dsDNA Abs were increased in *Rag^R229Q/R229Q^* mice. Thus, Inflammations in *Lig4^W447C/W447C^*mice are pathologically distinct from those in *Rag^R229Q/R229Q^*mice. The difference should come from broader expression of LIGIV than RAG. Non-lymphocytes with defective LIGIV, which are prone to DSB-induced effects such as cell death, should be involved in generation of inflammations.

Characterizing CD4^+^ TCR clonality has been used to monitor the progression of disease states in cases of autoimmunity ^32, 33^. TCR repertoires from *Lig4^W447C/W447C^* mice displayed increased clonality and decreased diversity compared with those from *Lig4^+/+^* mice. Among expanded T cell clones, activated Th1 cells were dominant, whereas *Il4*-or *Il17a*-expressing T cells were not observed. Further clarification of these T cell clones, which should be pathogenic, should contribute to effective treatment for the inflammations under acquired immunodeficiency.

In *Rag^R229Q/R229Q^* and *Lig4^R278H/R278H^* mice, splenic T cells showed oligoclonal expression of TCR Vβ gene-segments, but their usage of Vα was unclear ^19, 31^. In *Lig4^W447C/W447C^*mice, scRNA-seq analysis revealed that Vα and Jα gene-segments are used mainly towards the proximal locus, respectively. This skewed usage was not observed in Vβ and Jβ segments. This is consistent with usage pattern of TCRα gene segments not only from Omenn or Omenn-like syndrome patients ^34, 35^, but also from the mutant mice lacking XLF, which stimulates the DNA-joining activity of the XRCC4-LIGIV complex ^36^. It is interesting whether these skewed TCRαs are involved in intestinal inflammations.

In this study, mutant mice carrying a hypomorphic *Lig4* mutation derived from a LIG4 syndrome patient showed T cell-dependent intestinal inflammation under severe adaptive immunodeficiency. The inflammation depends on activation and/or selection of Th cells, especially Th1 cells. The phenotype is based on the monogenic mutation and can account for the inflammation accompanied with LIG4 syndrome, but the underlying pathological mechanisms should have some common aspects with polygenic intestinal bowel diseases, which are still intractable. Our mutant mice should be useful model mice to analyze and treat not only LIG4 syndrome itself, but also some common inflammatory conditions based on T cell activation and/or selection under adaptive immune deficiency.

## Methods

### *Lig4^W447C/W447C^* mice generation and genotyping

These *Lig4^W447C/W447C^* mice were generated using the CRISPR-Cas9 method ^37^. Briefly, guide RNA (gRNA; guide sequence: 5’-CAGACAAAAGAGGTGAAGGG-3’) and Cas9 endonuclease mRNA were generated *in vitro* using the MEGAshortscript T7 (Life Technologies) and mMESSAGE mMACHINE T7 ULTRA (Life Technologies) transcription kits, respectively. The synthesized gRNA and Cas9 mRNA were purified using the MEGAclear kit (Life Technologies). Single-stranded oligodeoxynucleotides containing the *Lig4* c.1341G>T mutation (5’-CCCCCCACAATTAGGACGTCTAATTCATCCATTAGTCCACTGACGTACTCTGGTTTAATCTTTAGACACCCTTCACCTCTTTTGTCTGGCTTGTAAATGGACAGAGGGTGTTTAACCATGATCCCCTCTTC-3’) were synthesized by Integrated DNA Technologies, Inc. To generate mutant mice, female B6C3F1 mice were super-ovulated and mated with C57BL/6 N males (CLEA Japan Inc.). Fertilized one-cell-stage embryos were injected with gRNA, Cas9 mRNA, and single-stranded oligodeoxynucleotides. The c.1341G>T mutation was confirmed by Sanger sequencing, and the mutant mice were further backcrossed to C57BL/6 N mice for more than six generations. We studied 8–16-week-old mutant mice and their littermates unless otherwise noted.

Genotyping was performed *via* polymerase chain reaction (PCR) amplification of tail DNAs using high-single nucleotide discrimination DNA polymerase (HiDi DNA polymerase; myPOLS Biotech GmbH). The following primers were used: control forward primer (CFP), 5’-AAGCCAGACAAAAGAGGTGAAGGGTGG-3’; mutation forward primer (MFP), 5’-AAGCCAGACAAAAGAGGTGAAGGGTGT-3’; and common reverse primer (CRP), 5’-TTTCATGGTGTAACCAGACCCAACACG-3’. PCR was performed with an initial denaturation at 95°C for 2 min, followed by 35 cycles of denaturation at 95°C for 15 seconds, annealing at 69°C for 10 seconds, extension at 72°C for 30 seconds, and then a final extension at 72°C for 1 min. The wild-type and mutant alleles were detected as a 225 bp band using PCR with CFP-CRP and MFP-CRP primer pairs, respectively.

### Generation of Ifng^−/−^, Lig4^W447C/W447C^Rag2^−/−^, and Lig4^W447C/W447C^ Ifng^−/−^ mice

*Ifng^−/−^* mice were generated using the Alt-R CRISPR-Cas9 System (Integrated DNA Technologies, Inc.). CRISPR RNAs (crRNAs) including the target sequence, gRNA1 (Exon 2; 5’-GTCTCTTCTTGGATATCTGGAGG-3’) or gRNA2 (between Exons 2 and 3; 5’-CATCAGCTGATAAAGCTAGGAGG-3’) were designed, annealed with tracrRNA, and then incubated with Cas9 protein to generate the gRNA-Cas9 complex according to the manufacturer’s instructions. The gRNA-Cas9 complex (the final 5 μM crRNA, 5 μM tracrRNA, and 1 μM Cas9) was injected into single-cell-stage fertilized eggs from C56BL/6N *via* electroporation (NEPA21 electroporator; Nepa Gene Co., Ltd). Approximately 350 injected eggs were transferred to the oviducts of pseudo-pregnant ICR females. After sequencing the PCR products of the targeted locus, six of the eight founders were found to be the *Ifng* variant. After that, one founder animal was selected to generate *Ifng* knockout mice.

Genotyping was performed *via* PCR amplification of tail DNAs using PrimeSTAR^®^ Max DNA Polymerase (TaKaRa). The following primers were used: forward primer, 5’-CTACGGTCAATCCTCTCCTCAC-3’; and reverse primer, 5’-TTTGGATTCTCACGGCCATAC-3’. PCR was performed with an initial denaturation at 98°C for 5 min, followed by 35 cycles of denaturation at 98°C for 10 seconds, annealing at 68°C for 5 seconds, extension at 72°C for 1 min, and then a final extension at 72°C for 1 min. The wild type was detected as a 1,620 bp band, while mutant alleles were detected as a 1224 bp band using PCR with a combination of primer pairs.

Heterozygous *Lig4^W447C/+^* mice were mated to *Rag2^−/−^*or *Ifng^−/−^* mice to generate *Lig4^W447C/W447C^ Rag2^−/−^* and *Lig4^W447C/W447C^ Ifng^−/−^* mice. C57BL/6N mice were used, and *Rag2* was genotyped using PCR, as previously described ^38^. We thank Dr. Frederick W. Alt for the *Rag2^−/−^* mice ^38^.

### Mice

C57BL/6N mice were purchased from CLEA Japan. All mice were housed in specific pathogen-free conditions with 12-hour light/dark cycles at an ambient temperature of 20-24°C and a humidity range of 40-60%. Experimental and control animals were co-housed and used at 8-16 weeks old unless otherwise noted. Both males and females were used in the study. Mice were euthanized by cervical dislocation before tissue dissection. All animal experiments were approved by the Animal Research Committee of Wakayama Medical University (Approval number 855 and 1140) and conducted in accordance with the Guidelines for the Care and Use of Laboratory Animals at Wakayama Medical University, as well as the relevant national guidelines and regulations.

### Preparation of MEFs

E14.5 embryos were removed, one by one, from the uterine, and the head-removed bodies were cut into small pieces with scissors. Then, they were cultured in a T-25 (25 cm^2^) culture flask (Falcon; Corning Inc.) per embryo with 2 ml of αMEM (Nacalai Tesque) supplemented with 10% fetal bovine serum (Gibco; Thermo Fisher Scientific) at 37[ under humidified 5% CO_2_ conditions. After two-day culture, the cells were fed with 2 ml of the medium, and then the medium was changed at another two-day culture. When growing cells became confluent, they were suspended in the medium supplemented with 10% dimethyl sulfoxide (Nacalai Tesque) and frozen in liquid nitrogen for stock. The frozen stock of MEFs were thawed and allowed to grow for subsequent experiments.

### **γ**H2AX foci assay

MEFs were plated at a density of 1×10^5^ on coverslips and incubated overnight. Then, the cells were irradiated with 1 Gy of X-rays by an X-ray machine (OM-B205; Ohmic) operated at 70 kVp and 5 mA using a 0.5 mm Al filter at room temperature. The dose rate was 0.64 Gy/min. After X-irradiation, the cells were incubated at 37[ for 10 min, 0.5, 1, 3, 6, and 24 h, then fixed with 4% paraformaldehyde phosphate buffer (Nacalai Tesque) for 15 min, and permeabilized with 0.5% triton X-100 (FUJIFILM Wako) in PBS (Gibco, Thermo Fisher Scientific) (-) on ice for 5 min. The fixed cells were soaked with blocking buffer containing 5% Blocking One (Nacalai Tesque) and 0.1 % Tween 20 (Sigma-Aldrich) in PBS (-) for 1 h. The blocked cells were incubated with an anti-phospho-histone H2AX (Ser139) mouse monoclonal antibody conjugated with Alexa Flour 488 (BioLegend) for 1 h at room temperature. After washing with 0.1 % Tween 20, the cells were counterstained with 4’,6-diamidino-2-phenylindole (DAPI; Sigma-Aldrich) and coverslips were placed on slide glasses followed by sealing with clear nail polish. For counting the number of γ-H2AX foci accurately and reproducibly, a newly developed Image J-based computer program was used as previously described ^39^.

### Flow cytometric analysis

A single-cell suspension was prepared from the thymus, spleen, BM, MLNs, and LP cells and analyzed using flow cytometry according to standard protocols. The LIVE/DEAD Fixable Aqua Dead Cell Stain Kit (Invitrogen) was used to exclude dead cells. The anti-CD16/32 antibody (2.4G2; Tonbo Biosciences) was used to block Fc receptors, and the cells were then stained with fluorochrome-conjugated or biotinylated antibodies **(Supplementary Table 5)**. For intracellular staining of IFN-γ, IL-17A and 1L-4, splenocytes, MLN cells, and LP cells were stimulated with PMA (50 ng/mL, Nacalai Tesque), ionomycin (500 ng/mL, Nacalai Tesque) and Golgi plug (BD Biosciences) for 5 hours. Cells were stained with monoclonal antibodies for CD3ε, CD4, and CD8α, then fixed with fixation/permeabilization solution (Cytofix/Cytoperm Kit, BD Biosciences), and stained with anti-IFN-γ, anti-IL-17A or anti-IL-4 monoclonal antibodies **(Supplementary Table 5)**. Foxp3 Staining Buffer Set (eBioscience) was used to stain Foxp3. Stained cells were analyzed using a FACSVerse flow cytometer (BD Biosciences), and data were analyzed using the FlowJo software Version 10.8.0 (BD Biosciences).

### Histological examination

The thymus, spleen, small intestine, and colon were formalin-fixed, paraffin-embedded, processed, and sectioned into 5-μm-thick sections. In addition, the BM was pre-decalcified. The sections were stained with hematoxylin and eosin. Colitis and enteritis were histo-morphologically assessed. Colitis was assessed using the Mouse Colitis Histology Index (MCHI) rating scale, as previously described by Koelink et al. ^40^. The total MCHI score ranged from 0 (no disease) to 22 (severe disease) and was calculated using four categories (goblet cell loss, crypt density, hyperplasia, and submucosal invasion) as follows: MCHI = 1 × goblet cell loss (0 to 3) + 2 × goblet density (0 to 2) + 2 × hyperplasia (0 to 3) + 3 × submucosal invasion (0 to 3). Enteritis was assessed using different criteria, as described in Supplementary Methods. Furthermore, we performed immunohistochemical analysis as previously described ^41^. Briefly, the thymus, spleen, small intestine, and colon were formalin-fixed, paraffin-embedded, and processed to obtain 5-μm-thick sections. In addition, we performed antigen retrieval using a microwave and a citrate buffer (pH 6.0; Sigma-Aldrich). Then, tissue samples were incubated at room temperature for 60 min with primary antibodies. The following primary antibodies were used: anti-mouse CD3 antibody (CD3-12; abcam), mouse anti-mouse CD4 antibody (EPR19514; abcam), anti-mouse F4/80 antibody (D2S9R; Cell Signaling Technology), and anti-mouse B220 antibody (RA3-6B2; BD). Next, DAB (3,3’-diaminobenzidine; Sigma-Aldrich) staining was performed using the two-step EnVision+ System-HRP methodology (Dako, Tokyo, Japan). For visualization, light microscopy was performed using an Olympus BX43 microscope (Olympus). Virtual slides were generated using the NanoZoomer S60 C13210 (Hamamatsu Photonics), and digitized whole-slide images (WSIs) were acquired. Images for analysis were subsequently extracted using NDP.view2Plus software (Hamamatsu Photonics).

### Western blot analysis

In brief, MEFs from *Lig4*-mutated mice were lysed with radio-immunoprecipitation assay lysis buffer (Nacalai Tesque) and cOmplete™ Mini Protease Inhibitor Cocktail (Roche Diagnostics). The protein extracts were mixed with Laemmli sample buffer containing 2-mercaptoethanol (Wako) and heated in a heat block at 100°C for 5 min. The samples were separated using sodium dodecyl sulfate-polyacrylamide gel electrophoresis and transferred onto polyvinylidene fluoride membranes (GE Healthcare). The membranes were washed and blocked with the DIG Wash and Block Buffer Set (Roche Diagnostics) according to the manufacturer’s instructions. The membranes were then incubated overnight at 4°C with primary antibodies. The following primary antibodies were used: rabbit anti-LIG4 (cat# ab26039; Abcam), mouse anti-XRCC4 (cat# MA5-24383; Thermo Fisher Scientific), rabbit anti-XLF (cat# ab33499; Abcam) and mouse anti-β-actin antibodies (cat# 3700, Cell Signaling Technology). The following day, after three washes, the membranes were incubated with secondary antibodies at room temperature for one hour. The following secondary antibodies were used: horseradish peroxidase-conjugated goat anti-rabbit IgG polyclonal antibody (cat# 170-6515, 1:1000 dilution; Bio-Rad) or horseradish peroxidase-conjugated rabbit anti-mouse IgG antibody (cat# 1080-05, 1:1000 dilution; SouthernBiotech). The protein bands were visualized with SuperSignal^®^ West Dura Substrate (Thermo Fisher Scientific) and detected using a LuminoGraph I Chemiluminescent Imaging System (ATTO). We measured the band intensities using Image J software version 1.51k (National Institutes of Health). The β-actin level was used as an internal control to ensure equal amounts of loading proteins.

### Measurement of serum Ig levels in mice

Serum Ig levels of *Lig4^W447C/W447C^* mice and wild-type C57BL/6N littermates were measured using an enzyme-linked immunosorbent assay (ELISA), as previously described ^37^. In brief, serum samples were incubated in an ELISA plate (Thermo Fisher Scientific) coated with anti-mouse IgM, IgG1, IgA or IgE antibodies and detected with biotinylated antibodies against each class and streptavidin-conjugated alkaline phosphatase (Southern Biotech). The plate was then developed using an alkaline phosphatase buffer (NaHCO_3_ [50 mM], MgCl2 [10 mM], pH 9.8) containing a phosphatase substrate tablet (Sigma-Aldrich). Purified mouse IgM, IgG1, IgA or IgE were used as standards. Details of all antibodies and standards are listed **(Supplementary Table 6)**.

### Mouse BM transplantation

BM cells were isolated from the femurs and tibias of *Lig4^W447C/W447C^* mice and wild-type C57BL/6N littermates (9–22 weeks of age). BM cells (0.5 to 1.0 × 10^7^) were injected *via* the tail vein into *Rag2^−/−^* recipient mice (10–30 weeks of age) that had received a lethal total-body γ-irradiation dose of 10 Gy. Mice were monitored daily for survival and weighed twice a week. Euthanasia was performed seven weeks after transplantation for analysis.

### Bulk RNA-Sequencing analysis

For RNA-sequencing (RNA-Seq) analysis, total RNA was extracted using the RNeasy Micro Kit (QIAGEN) or the Sepasol-RNA I Super G (Nacalai Tesque), as previously described ^42, 43^. The samples were collected from the colon, MLNs and spleen of *Lig4^+/+^* and *Lig4^W447C/W44^*^7^ and from the colon of *Lig4^W447C/W447C^Ifng^−/−^*mice. The RNA quality was assessed using an Agilent 4200 TapeStation (Agilent Technologies), and the RNA concentration was measured using a Qubit Fluorometer (Thermo Fisher Scientific). A total of 4,000 ng RNA in colon, 228–1,552 ng RNA in MLN and 921.6–3,000 ng RNA in spleen was used, and libraries for sequencing were constructed using TruSeq Stranded mRNA (Illumina Inc.) according to the manufacturer’s protocol. The concentration of libraries was estimated using the KAPA Library Quantification Kit (Roche Diagnostics). The range of average library size was 324–359 bp. The libraries were sequenced by high-throughput sequencing using a NextSeq 500/550 High Output Kit v2.5 (Illumina, 75 cycles pair-end, 40/40 cycles). The average number of sequence reads per sample was 22,855,222.1. The bulk RNA-seq results were analyzed using the CLC Genomics Workbench Version 12.0.2 (Filgen Inc.), and the abundance of gene expression was expressed using the reads per kilobase per million values. In addition, gene expression analysis, including gene set enrichment analysis (GSEA), was conducted using GSEA v4.2.3 software and the Molecular Signature Database v7.5.1.^44, 45^. Moreover, the web tool ClustVis and BioJupies were used to draw a heatmap and a volcano plot, respectively ^46, 47^. The raw fastq data of RNA-seq analysis was deposited in DDBJ and BioProject number is PRJDB19195, Run number is DRR615557-DRR615579.

### Single-cell RNA-sequencing and TCR repertoire analysis

Single-cell transcriptome and TCR repertoire analysis were performed using Chromium Single Cell 5’ Reagent Kits v2 and Chromium Single Cell Human TCR Amplification Kits with Chromium Controller (10x Genomics) according to the manufacturer’s instructions as previously described ^48^. Libraries were sequenced on NovaSeq X Plus (Illumina) as paired-end mode (read 1: 151 bp; read 2: 151 bp). The raw reads were processed by Cell Ranger v7.1.0 (10x Genomics). Gene expression– based clustering was performed using the Seurat R package (v3.1). Cells with a mitochondrial content >10% and cells with <200 or >4,000 genes detected were considered outliers (dying cells and empty droplets and doublets, respectively) and filtered out. The Seurat SCTransform function was used for normalization, and data were integrated without performing batch-effect correction as all samples were processed simultaneously. Hashtag oligo demultiplexing was performed on centered log ratio-normalized hashtag unique molecular identifier counts, and clonotypes were matched to the gene expression data through their droplet barcodes, using Python scripts. The cluster analysis based on t-SNE plot and differential expression analysis were performed by Loupe Browser v8.0.0 (10x Genomics). The TCR repertoire analysis was performed by Loupe V(D)J Browser v5.2.0(10x Genomics). The single-cell RNA-seq data generated in this study have been deposited in GEO under accession number GSE294169.

### LIGIV protein structure predictive analysis

Atomic models of wild-type and mutated LIGIV proteins were predicted with AlphaFold3 webserver with default settings ^49^. The amino acid sequence of mouse LIGIV was used as the input. Five models were generated and the relaxed model with the highest confidence score was selected for analysis. The molecular graphics were made using PYMOL (http://www.pymol.org/).

### Cell preparations

Thymocyte, splenocyte, and MLN suspensions were prepared by grinding the appropriate organs through mesh filters. In addition, BM cells were extracted from the femur and tibia and passed through mesh filters. Furthermore, intestinal immune cells were prepared from the small intestine. In brief, fat and Peyer’s patches were dissected away before the small intestine was opened longitudinally and stirred in RPMI 1640 media (Nacalai Tesque) containing 2% fetal bovine serum (JR Scientific, Inc.) and 2 mM EDTA (Nacalai Tesque) for 20 min at 37°C. IELs were isolated from the supernatant using a 40%–75% Percoll density gradient (GE Healthcare) by collecting cells that layered between the 40% and 75% fractions. Following supernatant collection, the intestinal tissue was stirred for 20 min at 37°C in RPMI 1640 media (Nacalai Tasque) containing 2% fetal bovine serum before mincing and further stirring in 400 units/mL of collagenase D (Roche Diagnostics) and 10 μg/mL of DNase I (Sigma-Aldrich) for 20 min at 37°C. Floating cells were collected, and the collagenase digestion was repeated twice more. The pooled cell suspensions were then centrifuged on a 40%–70% Percoll density gradient, and cells that layered between the 40%–75% fractions were collected as LP cells.

### Statistical analysis

Statistical analysis was performed using GraphPad Prism 9 (GraphPad Software). The data were compared using an unpaired student’s *t*-test, unless otherwise specified, and presented as the mean and standard error of the mean (SEM). *P*-values < 0.05 were considered statistically significant. Unless otherwise noted, \**p* < 0.05, \*\**p* < 0.01, and \*\*\**p* < 0.001. N.S., non-significant differences.

## Supplementary Methods

### Measurement of serum anti-dsDNA antibodies levels in mice

Serum anti-dsDNA antibodies levels of *Lig4^W447C/W447C^* mice and wild-type C57BL/6N were measured by enzyme-linked immunosorbent assay (ELISA) using LBIS Mouse anti-dsDNA ELISA Kit (Cat# 637-02691, FUJIFILM), following the manufacturer’s instructions.

### Histopathological evaluation of enteritis

Enteritis was assessed using the inflammatory score reported by Erben et al.^1^. The inflammation score ranged from 0 (no disease) to 5 (severe disease) and was determined based on the severity and extent of inflammatory cell infiltration, epithelial changes, and mucosal architecture.

### Gut microbiota analyzed by *16S rRNA* gene sequencing

Stool samples were collected and sent to ICLAS Monitoring Center, Central Institute for Experimental Animals for gut microbiota analysis. The samples were homogenized, and part of the homogenate was suspended in a stool collection kit (TechnoSuruga Lab). DNA was extracted from the samples using the beads-phenol method. The extracted DNA was then used for terminal restriction fragment length polymorphism analysis targeting bacterial 16S rDNA. For fragment analysis, an ABI PRISM 310 Genetic Analyzer (Applied Biosystems) and GeneScan software (Applied Biosystems) were used. The length of each fragment was determined based on operational taxonomic unit, and the major bacterial taxa were roughly estimated using an in-house mouse gut microbiota database constructed by the laboratory.

## Acknowledgments

This work was supported by Grant-in-Aid for Transformative Research Areas (JP22H05182 and JP22H05187 to T. Kaisho. and JP22H05187 to I.S.), for Scientific Research (B) (JP26293106, JP17H04088, JP20H03505 and JP24K02298 to T. Kaisho), for Scientific Research (C) (JP19K07628 and JP22K07006 to I.S., JP21K12243 to S.K., JP19K08821 to T.S., JP20K08718 and JP23K07865 to S.T.), for Scientific Research on Innovative Areas (JP17H05799 and JP19H04813 to T. Kaisho), for Early-Career Scientists (JP20K17405 and JP22K16308 to Y.Y., JP25K19582 to H.K.), for Research Activity Start-up JP23K19487 to T.Kato), for JSPS Fellows (JP21J22615 to T. Kato), for Exploratory Research (JP17K19568 and JP21K19384 and JP23K18222 to T. Kaisho) from the Japan Society for the Promotion of Science, Takeda Science Foundation (to Y.Y., I.S. H.H. and T. Kaisho), GSK Japan Research Grant 2021 (to I.S.), Kowa Life Science Foundation (to I.S.). This work was also supported in part by the Wakayama Medical University Special Grant-in-Aid for Research Projects (K23TS03 to S.T.). We acknowledge proofreading and editing by Benjamin Phillis at the Clinical Study Support Center at Wakayama Medical University. We thank the Laboratory Animal Center at Wakayama Medical University for their technical support in generating the *Ifng* knockout mice and Yukako Hirato at Osaka Metropolitan University for technical assistance of γH2AX foci assay.

## Author contributions

Y.Y., S. Tamura and T. Kaisho designed the research. Y.Y., H.K., T. Kato, I.S., T.O., M.T., S. Tabata, K.N., K.T., K.S., Y.U., A.S., S.K., K.O. and Y.F. performed biochemical and histochemical experiments. T.M. performed *in silico* analyses on the LIGIV protein structures. Y.Y., H.H., T. Kaisho and S. Tamura generated and analyzed murine models. Y.Y., H.K., S.I., D.O. and S.H. performed genomic analyses. Y.Y., N.K., T.S., T. Kaisho and S. Tamura wrote and edited the manuscript.

## Supplementary information

The online version contains Supplementary material available at….

**Correspondence and requests** for materials should be addressed to Yusuke Yamashita, Tsuneyasu Kaisho, and Shinobu Tamura.

## Extended Data Figures

**Extended Data Figure 1. Appearance and body weight of newborn mice, and** γ**H2AX foci assay in MEFs.** (a) The appearance of one-day-old *Lig4^+/W447C^* mice and *Lig4^W447C/W447C^*littermates. The number indicates units of centimeters. The graph (right) depicts the weight of one-day-old littermate mice. (*Lig4^+/ W447C^* [*n* = 4] and *Lig4^W447C/W447C^* [*n* = 3]) (b) γH2AX foci analysis in plateau-phase *Lig4^+/+^*, *Lig4^+/W447C^* and *Lig4^W447C/W447C^* MEFs after a 1 Gy irradiation. The data are presented as the mean ± standard error of the mean (SEM). *P*-values were calculated as follows: Within each MEF genotype, one-way analysis of variance (ANOVA) was conducted, followed by Tukey’s multiple comparison test. The Mann-Whitney U test was used to compare baseline measurements among different genotypes.

**Extended Data Figure 2. LIGIV protein structure predictive analysis**. Structural insights into R278H (a) and Y288C (b) mutations. (a) R278 (LIGIV wild-type) indicated cyan ribbon and LIGIV H278 did magenta ribbon. (b) Y288 (LIGIV wild type) indicated cyan ribbon and LIGIV C288 indicated yellow ribbon. (a and b, left panels) Neither mutation caused overall structural change. (a, right panel) R278 and H278 binding to AMP are shown in stick representation (cyan and magenta, respectively). R278H mutation decreased the interface area of AMP-binding. (b, right panel) R278 and H278 binding to AMP are shown in stick representation (cyan and yellow, respectively). Neither Y288 nor C288 interacted with AMP-binding.

**Extended Data Figure 3. Anti-dsDNA antibody levels in serum measured by ELISA**. Serum anti-dsDNA antibody levels in *Lig4^+/+^* and *Lig4^W447C/W447C^* mice (*Lig4^+/+^* [*n* = 12] and *Lig4^W447C/W447C^* [*n* = 12]; 9–23 weeks).

**Extended Data Figure 4. FACS analysis of thymic T cells and splenic regulatory T cells**. (a) The percentages dynamics of thymic T cells at various developmental stages, including double negative, double positive, and single positive (SP) in *Lig4^+/+^* and *Lig4^W447C/W447C^* mice (*Lig4^+/+^* [n = 4] and *Lig4^W447C/W447C^* [n = 4]; 7–16 weeks). (b) FACS analysis of splenocytes labeled with CD4, CD25 and Foxp3 antibodies from *Lig4^+/+^*and *Lig4^W447C/W447C^* mice. (c) Percentage of regulatory T cells in CD4+ splenocytes of *Lig4^+/+^* and *Lig4^W447C/W447C^* mice (*Lig4^+/+^* [*n* = 5] and *Lig4^W447C/W447C^* [*n* = 5]; 18–24 weeks) using FACS analysis.

**Extended Data Figure 5. Inflammation of the small intestine in *Lig4^W447C/W447C^* mice**. (a) Hematoxylin and eosin (H&E) staining and immunohistochemical staining with CD4, F4/80 and B220 antibodies of the small intestine from *Lig4^+/+^* and *Lig4^W447C/W447C^* mice. Scale bar represents 60 μm. (b) A small intestine inflammatory score of each mouse (*Lig4^+/+^* [*n* = 11] and *Lig4^W447C/W447C^* [*n* = 11]; 7–18 weeks). (c) FACS analysis of T cells labeled with T-cell receptor (TCR)β, TCRγδ, CD4, and CD8 antibodies in the lamina propria of *Lig4*^+/+^ and *Lig4^W447C/W447C^* mice. (d) The percentages of TCRβ^+^, TCRγδ^+^, and CD4^+^ T cells in the lamina propria of each mouse (*Lig4*^+/+^ [*n* = 7] and *Lig4^W447C/W447C^* [*n* = 6]; 9–12 weeks).

**Extended Data Figure 6. Gut microbiota analyzed by *16S rRNA* gene sequencing on the feces of *Lig4^+/+^* and *Lig4^W447C/W447C^* mice**. The proportion of intestinal flora did not differ significantly between *Lig4^+/+^*(*n* = 8) and *Lig4^W447C/W447C^* mice (*n* = 8). The data are presented as the mean ± standard error of the mean (SEM). *P*-values were calculated using the unpaired student’s *t*-test.

**Extended Data Figure 7. Intracellular cytokine staining I**ntracellular cytokine staining (IFN-γ, IL-17A, and IL-4) of CD4^+^ T cells in splenocytes from *Lig4^+/+^ Ifng*^+/+^, *Lig4^W447C/W447^*^C^*Ifng*^+/+^ and *Lig4^W447C/W447C^Ifng^−/−^* mice.

**Extended Data Figure 8. TCR repertoire analysis using bulk RNA-sequencing** (a-d) The bar graphs show the relative expression levels of each *Trav* and *Trbv* gene segment analyzed by bulk RNA-sequencing. Data are presented as a percentage of the total *Trav* and *Trbv* gene expression in individual *Lig4^+/+^* mice [n=3; White bars] and *Lig4^W447C/W447C^*mice [n=3; red bars]. Gene segments are arranged in chromosomal order, from the 5’ end (left) to the 3’ end (right) of the *Trav* and *Trbv* loci. (a) *Trav* expression in the spleen. (b) *Trav* expression in the MLN. (c) *Trbv* expression in the spleen. (d) *Trbv* expression in the MLN.

**Extended Data Figure 9. TCR repertoire heatmaps of splenocytes samples analyzed by single-cell RNA-sequencing**. Heatmaps showing the usage of *Trav*–*Traj* (a) and *Trbv*–*Trbj* (b) combinations. Splenocytes were prepared from individual mice (*Lig4^+/+^*[n=1] and *Lig4^W447C/W447C^* [n=3; exp #1, #2 and #3]) and analyzed by single-cell RNA-sequencing. (a) The x– and y-axis represents *Trav* and *Traj* segments, respectively, arranged from 5′ to 3′ end of the locus. The color gradient ranges from red to white, indicating high to low frequency of each *Trav*–*Traj* pair, respectively. The scale is shown on the right. (b) the x– and y-axis shows *Trbv* and *Trbj* segments, respectively, arranged from 5′ to 3′ end of the locus. The same color gradient as shown in (a) is used, with the scale shown on the right.

