## Extended Data Fig. 1-9 and Supplemental Table 1-6 for "Hypomorphic *Lig4* gene mutation in mice predisposes to Th1-skewing intestinal inflammation": Extneded Data Fig.1-9.pdf

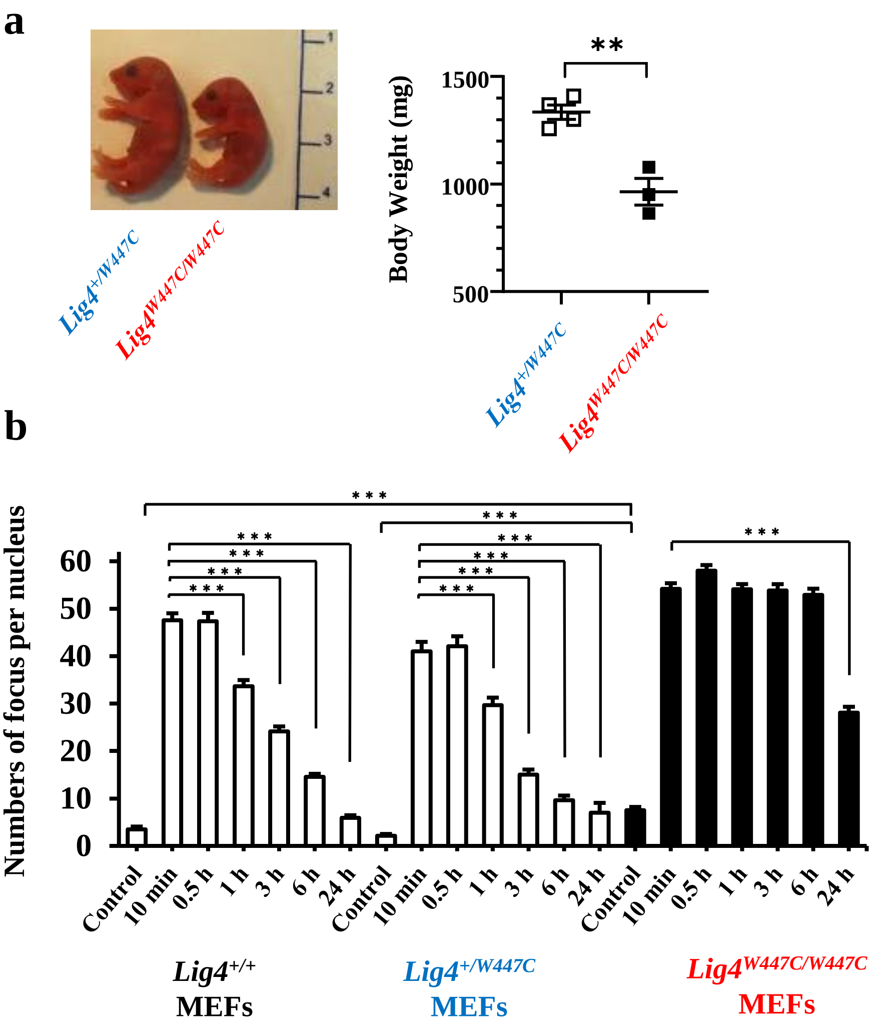


### Extended Data Fig. 1

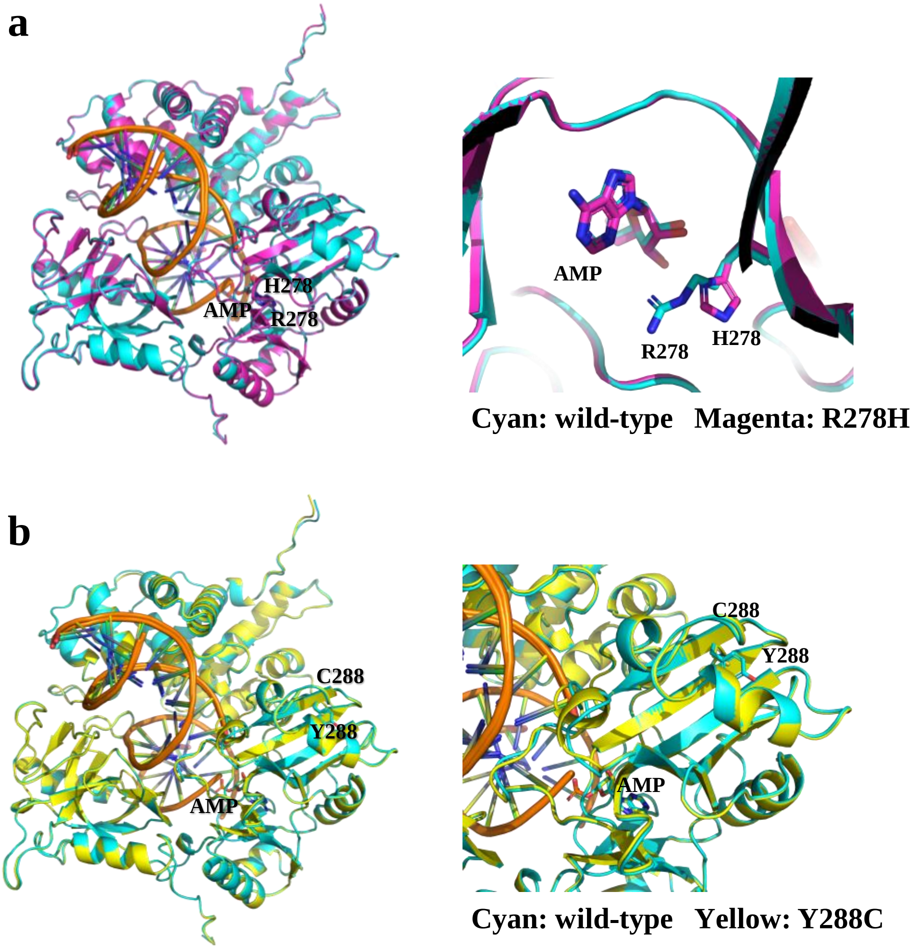


### Extended Data Fig. 2

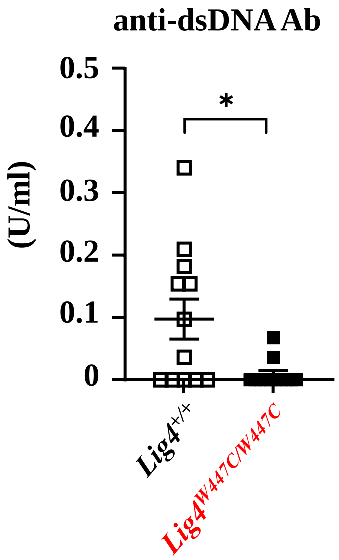


### Extended Data Fig. 3

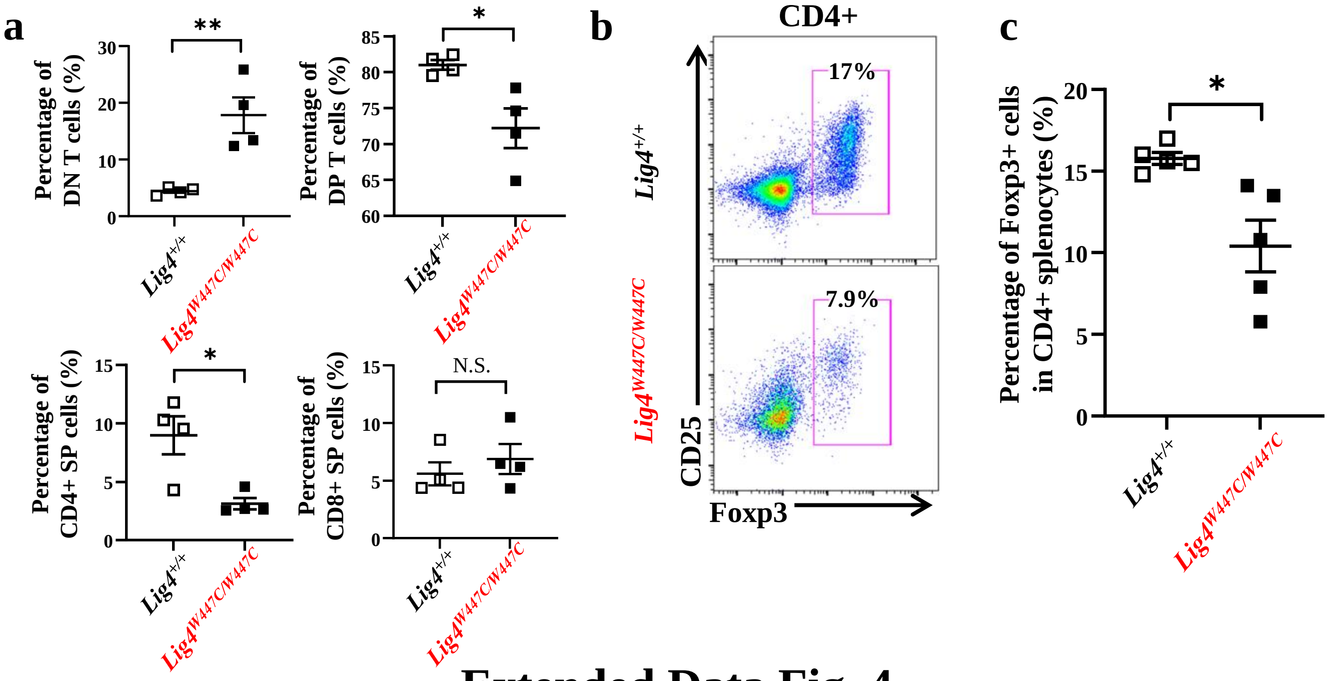


### Extended Data Fig. 4

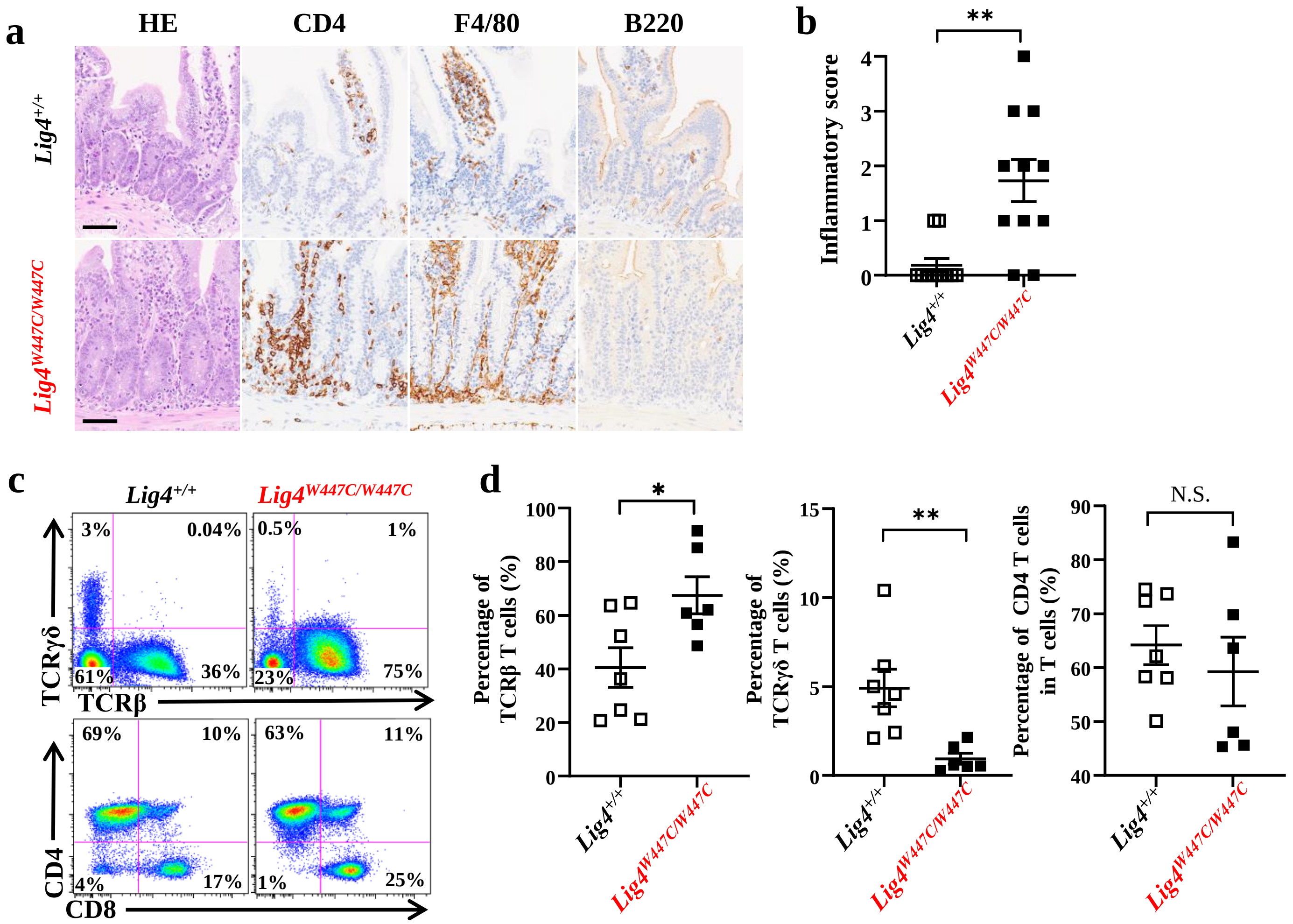


### Extended Data Fig. 5

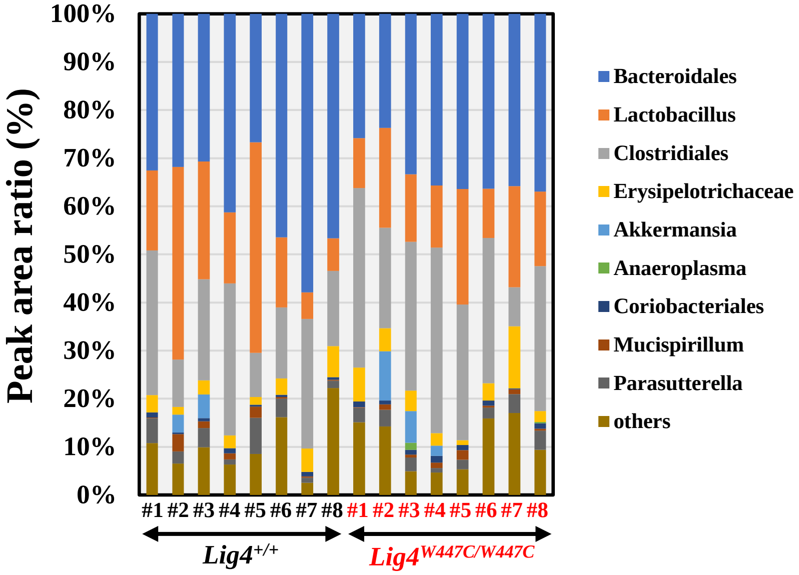


| Predicted bacteria | *Lig4*$^{+/+}$ | *Lig4*$^{W447C/W447C}$ | P value |
|---|---|---|---|
| Bacteroidales | 39.3±3.8 | 32.8±1.9 | 0.15 |
| Lactobacillus | 20.8±5.1 | 16.0±1.9 | 0.39 |
| Clostridiales | 19.9±3.1 | 27.8±3.4 | 0.11 |
| Erysipelotrichaceae | 3.4±0.6 | 4.8±1.3 | 0.35 |
| Akkermansia | 1.1±0.7 | 2.3±1.4 | 0.43 |
| Anaeroplasma | 0±0 | 0.2±0.2 | 0.24 |
| Coriobacteriales | 0.7±0.1 | 1.0±0.1 | 0.10 |
| Mucispirillum | 1.2±0.4 | 0.8±0.5 | 0.49 |
| Parasutterella | 3.3±0.8 | 2.8±0.4 | 0.53 |
| others | 10.3±2.2 | 10.8±1.9 | 0.89 |

### Extended Data Fig. 6

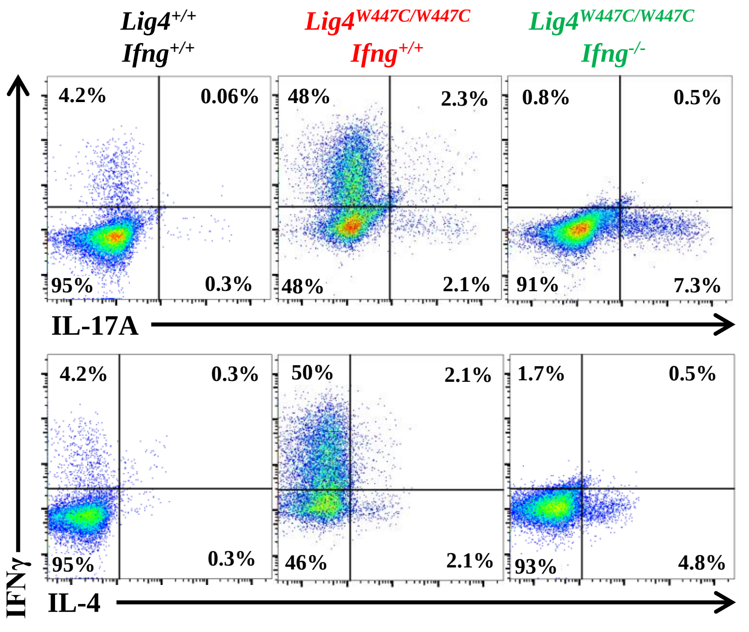


### Extended Data Fig. 7

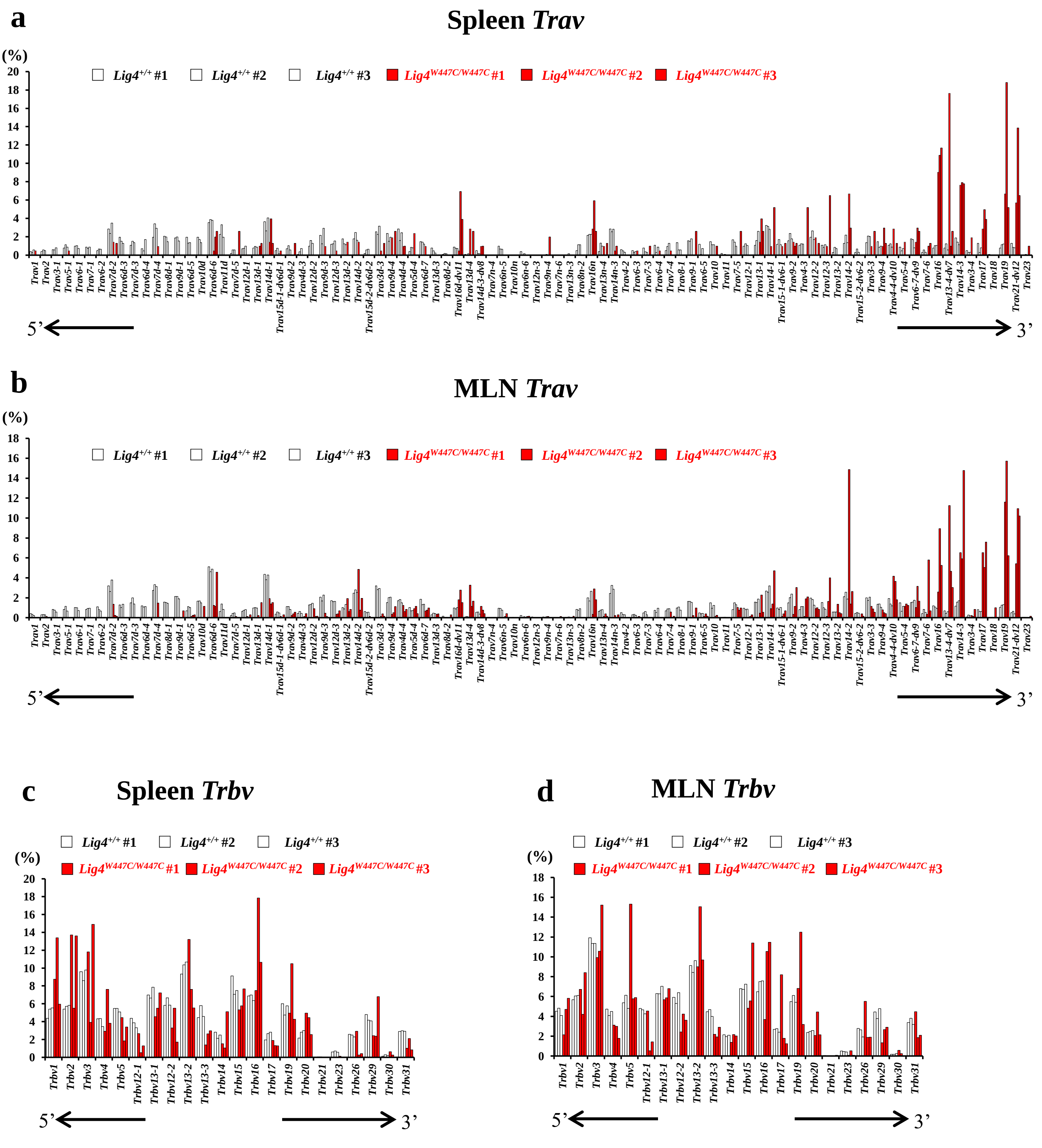


### Extended Data Fig. 8

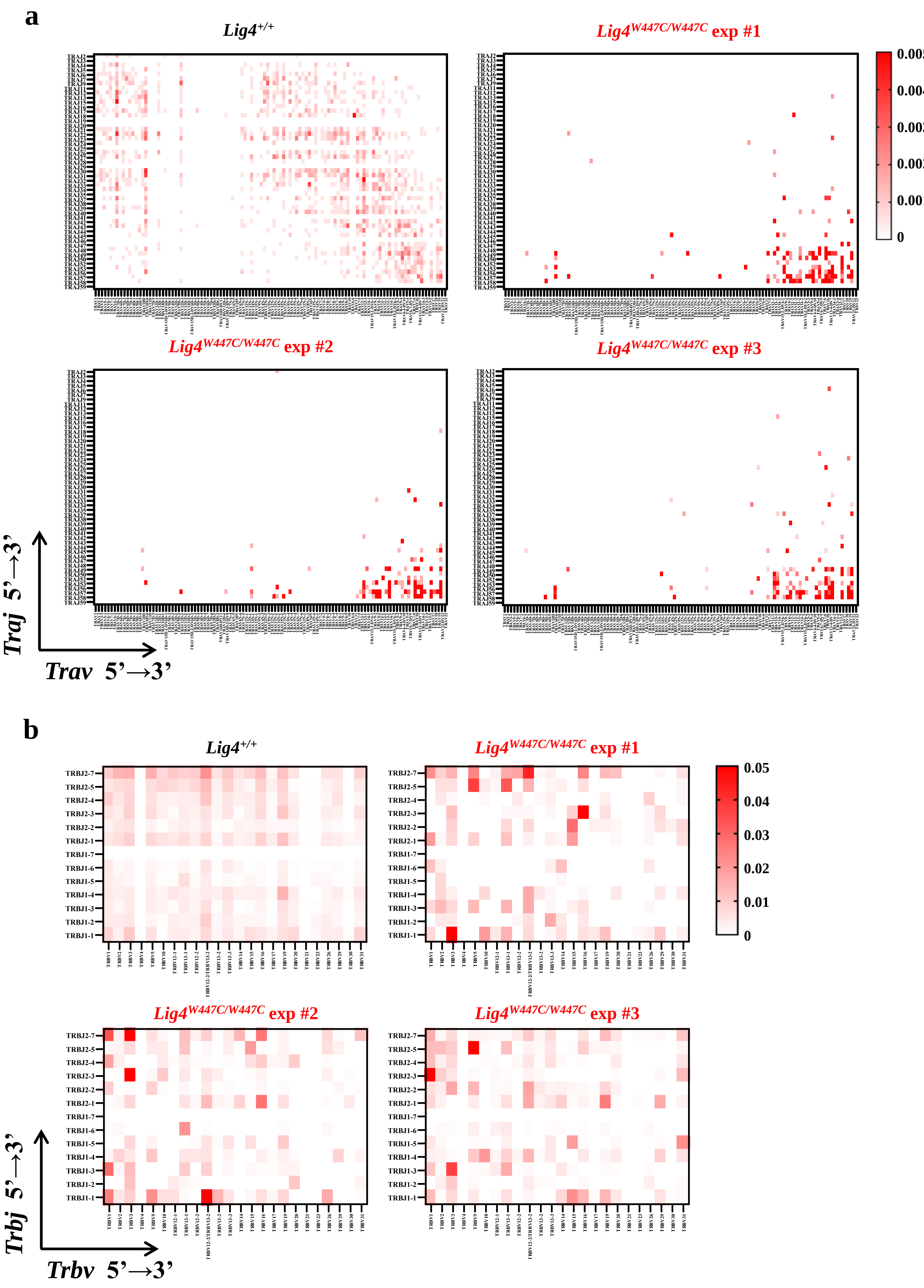


### Extended Data Fig. 9
